# Mechanism of counterattack against eIF2α kinase immune signalling by viral pseudo-kinase PK2

**DOI:** 10.1101/2025.02.05.636571

**Authors:** Daijiro Takeshita, Yuko Takagi

## Abstract

Viral infection triggers the activation of host innate immune responses, including the integrated stress response (ISR). The antiviral response is mediated by the phosphorylation of eukaryotic translation initiation factor 2 alpha (eIF2α) catalyzed by host eIF2α kinase, resulting in suppression of global protein synthesis and induction of selected genes in the cell. To evade and/or subvert the host antiviral signalling pathway, viruses have evolved and harnessed elaborate counter-attack mechanisms to repress the activation of host eIF2α kinase. In insects, eIF2α kinase-mimic protein PK2 encoded in baculovirus directly binds to and inhibits host eIF2α kinase for viral propagation. However, the mechanism underlying how viral PK2 recognizes host eIF2α kinase and interrupts its activity has remained unclear. Here we present a series of crystal structures of apo PK2 and its complex with eIF2α kinase, revealing a conformational transition mechanism of PK2 for eIF2α kinase inhibition. PK2 alone, comprised of an N-terminal extension (NTE) and eIF2α kinase C-lobe mimic (EKCM), adopted a two-fold symmetric homotetramer assembled mainly by the NTE region. Pull-down assay identified a region of *Bombyx mori* HRI-like kinase (BmHRI) that binds to PK2, and its complex structure revealed that the PK2-binding region of BmHRI, subdomain III-IV, is entrapped in a newly formed groove between the NTE region and EKCM of PK2. The structural data, together with biochemical analyses, indicated that PK2 suppresses eIF2α kinase activity by the pullout and subsequent blockade of the subdomain III-IV, a regulatory element essential for its kinase activity. These results not only provide the molecular mechanisms for the inhibition of host eIF2α kinase by viral pseudo-kinase, but also unveil the viral evolutionary strategy to shut-off host immunity via horizontal gene transfer from host to virus.

## Introduction

In nature, viruses are numerous and diverse, and evolve at a rapid rate to adapt to their hosts. Viruses rely on host cellular machineries for their replication and propagation, thus they have acquired and developed sophisticated strategies to evade and/or subvert host’s immune system for successful infection. Viral infection induces a series of antiviral innate immune responses initiated by host pattern-recognition receptors (PRRs), which recognize virus-associated molecules, such as genomic DNA, RNA or double-stranded RNA (dsRNA) (1, 2). Upon viral infection, nucleic acids or viral proteins generated by viral replication activates the integrated stress response (ISR) through the phosphorylation of eIF2α by eIF2α kinases, which leads to suppression of global protein synthesis and induction of selected genes, including activating transcriptional factor 4 (ATF4), that together serve to maintain the host physiological homeostasis (3–8).

Translational initiation factor eIF2, consisting of α, β and γ subunits, forms a ternary complex with GTP and methionyl initiator tRNA (Met-tRNAi^Met^) to deliver the initiator tRNA to the 40S ribosomal subunit and participates in pre-initiation complex (PIC) for translation initiation. Recognition of an AUG start codon on mRNA leads to the hydrolysis of GTP bound to eIF2, and then eIF2-GDP is released from PIC. Under normal conditions, the eIF2-GDP is recycled to the active eIF2-GTP by guanine nucleotide exchange factor (GEF) eIF2B for next round of translation initiation. However, the phosphorylation of eIF2α on Ser51in the ISR converts the substrate into the competitive inhibitor of eIF2B, leading to the inhibition of translation initiation (7, 8). In mammalian cells, there are four eIF2α kinases responsible for the eIF2α phosphorylation, namely PKR (protein kinase double-stranded RNA-dependent), GCN2 (general control non-derepressible-2), PERK (protein kinase R-like endoplasmic reticulum kinase) and HRI (heme-regulated inhibitor). These four kinases have unique regulatory domains for sensing different stress stimuli, while sharing homologous catalytic domains for the phosphorylation of eIF2α (5, 8). These eIF2α kinases are activated by different environmental signals, dsRNA for PKR, amino acid starvation for GCN2, unfolded or misfolded proteins for PERK, and heme deficiency for HRI, and in certain cases, overlapping stimuli cause redundant activation of these kinases (5). Depending on the host cells and viral species, viral infection can activate any of the four eIF2α kinases, inducing the ISR signalling (5, 6, 8, 9).

Mammalian PKR is the well-studied eIF2α kinase and plays a crucial role in innate immune system in response to viral RNAs (10–13). PKR is composed of two N-terminal double-stranded RNA binding domains (dsRBDs) and a C-terminal kinase domain. Upon binding of viral dsRNA to the dsRBDs, PKR is activated through the homodimerization and subsequent autophosphorylation at multiple serine/threonine sites, which in turn phosphorylates Ser51 of eIF2α (4, 7). The phosphorylated eIF2α inhibits translation initiation, suppressing the viral replication and propagation. To evade and/or suppress the host cellular immune response, many viruses have evolved elaborate counter-attack mechanisms to override the PKR-mediated immune signalling (14–17). K3L encoded in vaccinia virus shares an amino acid sequence homologous to the N-terminal third of eIF2α and inhibits PKR in a competitive manner (18–23). Along with the structural similarity between K3L and eIF2α, K3L would act as a pseudo-substrate and competitive inhibitor of PKR (23). Notably, human PKR has evolved to evade the viral mimics by means of positive selection mostly around the binding site of eIF2α to maintain their primary function, suggesting molecular evolutionary ‘arms-races’ between PKR and the viral antagonist (24, 25). In addition, other types of eIF2α kinase, PERK, GCN2 and HRI have been shown to serve for the host immune signalling against viral infection, and therefore the antiviral immune response of eIF2α phosphorylation mediated by eIF2α kinases would be a common defense mechanism in host eukaryotic cells (8, 9, 17).

PK2 encoded by a baculovirus *Autographica californica* multiple nucleopolyhedrovirus (AcMNPV) is an anti-eIF2α kinase factor that is conserved among alphabaculoviruses. PK2 expressed in AcMNPV-infected cells inhibits eIF2α phosphorylation in cells, and interacts with human PKR *in vitro* and inhibit its kinase activity (26–29). In Sf9 cells, UV-induced eIF2α phosphorylation and caspase activation are reduced in wildtype baculovirus infection, but not in pk2-lacked baculovirus infection (28). The inhibition of eIF2α phosphorylation by PK2 is required for efficient viral infection in *Bombyx mori*, both in cultured cells and larval insects (29). In addition, PK2 overexpression in Sf9 cells improves the production of progeny virus and enhanced insecticidal activity against *Spodoptera exigua* larvae (30). Collectively, these results have shown that baculovirus PK2 is crucial for viral fitness via inhibition of the eIF2α kinase activity. PK2 is a pseudo-kinase, homologous to the C-lobe of eIF2α kinase, and the phylogenetic analysis showed that the PK2 gene originally evolved from an insect HRI-like kinase through horizontal gene transfer from host to viral genome (29). PK2 inhibits *B. mori* HRI-like kinase (BmHRI) more efficiently than other eIF2α kinases in vitro and in vivo, and knockdown of BmHRI in host cells can rescue viral production for a *pk2*-deficient virus, indicating that the HRI would be the natural target of PK2 (29).

The amino acid sequence of PK2 contains two distinct regions, a 22-residue <u>N</u>-terminal <u>e</u>xtension (NTE) and a C-terminal domain homologous to the protein kinase, named <u>e</u>IF2α <u>k</u>inase <u>C</u>-lobe <u>m</u>imic (EKCM) domain (27, 29, 31) (**Fig. 1a**), both of which are shown to be necessary for the eIF2α kinase inhibition (29). Pulldown assay and two hybrid assays showed that PK2 interacts with the N-lobes of PKR and BmHRI but not with the C-lobes, which suggest that PK2 displaces the C-lobe of the kinase domain. These results led to a “PK2 lobe-swapping inhibition model”, in which a nonfunctional pseudokinase-kinase complex, composed by the interaction between the kinase N-lobe and PK2, perturbs the ATP-binding and inhibits the substrate phosphorylation (29). However, due to the lack of the structural information of PK2 in complex with eIF2α kinase as well as its apo form, the precise molecular mechanism of host eIF2α kinase inhibition by PK2 has remained unclear.

**Figure 1.**
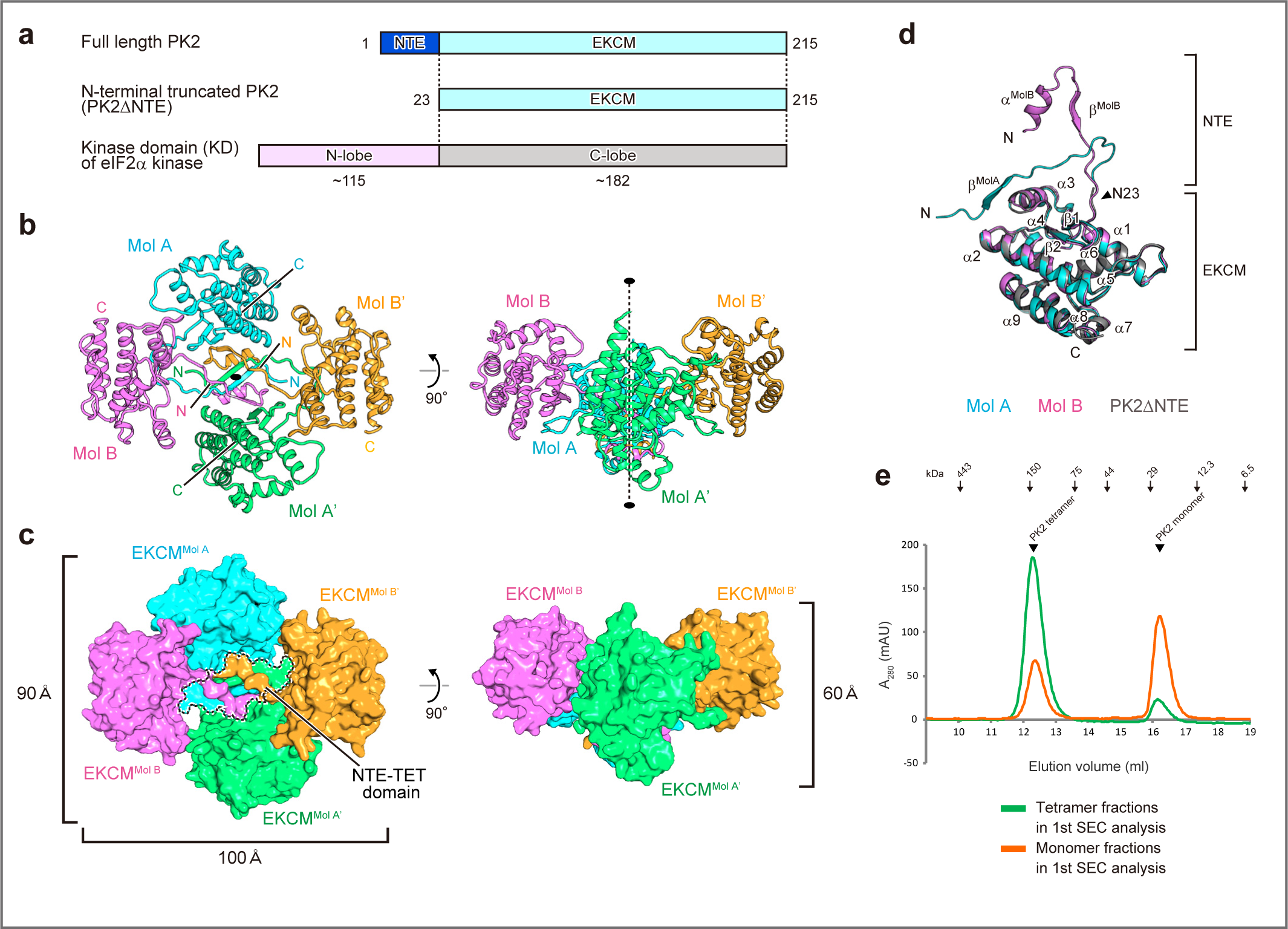
Overall structure of PK2. **(a)** Domain organization of PK2 composed of NTE region and EKCM. Full-length PK2 and PK2ΔNTE lacking the NTE region are shown in the upper and middle, respectively. For comparison, domain organization of the kinase domain of eIF2α kinase (human PKR^KD^) is shown in the bottom. **(b)** Ribbon representation of PK2. The four protomers are labeled Mol A, Mol B, Mol A’, and Mol B’ and colored cyan, magenta, green, and orange, respectively. Dashed line between ellipses indicates the non-crystallographic two-fold axis of the symmetry. **(c)** Surface representation of PK2 colored same as in Fig. 1b. NTE-TET domain is encircled by a black dashed line. **(d)** Superposition of PK2 protomers, Mol A (cyan) and Mol B (magenta), and PK2ΔNTE (gray). Stereoview of this Fig. 1d is shown in Extended Data Fig. 1f. **(e)** Representative gel filtration chromatograms of PK2 samples, including tetrameric fractions (green) and PK2 monomeric fractions (orange) of 1^st^ SEC analysis.

To elucidate the molecular mechanism of the eIF2α kinase inhibition by PK2, we firstly determined the crystal structures of the full-length PK2 and the truncated PK2, PK2ΔNTE, lacking the NTE region, at 2.7 Å and 2.8 Å resolutions, respectively. The crystal structure of the full-length PK2 revealed that the PK2 forms a tightly associated homotetramer mainly via the NTE region positioned at the center of the complex. The full-length PK2 exists in a monomer and homotetramer equilibrium in solution, whereas PK2ΔNTE exists as a monomer in both solution and crystal structure. We further determined the crystal structures of PK2 in complex with the eIF2α kinase PK2-binding region, named here as PK2-BR, at 2.0-2.1 Å resolution. The complex crystal structures revealed that PK2-BR, including a key regulator element αC helix and β4 strand of eIF2α kinase, intrudes into a cleft between the NTE region and EKCM of PK2. The complex structures explain how PK2 inhibits the eIF2α kinase activity through the specific interactions between PK2 and PK2-BR of BmHRI. Our comprehensive structural and biochemical studies elucidate the molecular and functional mechanisms underlying the inhibition of eIF2α kinase by pseudo-kinase PK2. Furthermore, these findings offer valuable insights into the viral evolutionary strategies to counteract host innate immune response through horizontal gene transfer.

## Results

### Structural determination of PK2 and PK2ΔNTE in the apo form

To elucidate the molecular mechanism of eIF2α kinase inhibition by PK2, we first set out to determine the crystal structure of baculovirus PK2 in the apo form. The full-length PK2 was crystallized (32) and its structure was determined using the single-wavelength anomalous dispersion (SAD) method. The structure of PK2 was refined to *R* factor of 23.7% (*R*_free_ = 29.9%) at 2.7 Å resolution (**Extended Data Table 1**). The asymmetric unit in the crystal contained four molecules of PK2, which form a tightly associated homotetramer. The N-terminally truncated PK2 protein, PK2ΔNTE, which contains only the EKCM domain (residues 23-215) lacking the NTE (Fig. 1a), was also crystallized (32) and its structure was determined by the molecular replacement. The structure of PK2ΔNTE was refined to 2.8 Å resolution with *R* factor of 22.7% (*R*_free_ = 28.2%) (**Extended Data Table 1**). Contrary to the full-length PK2 structure, the PK2ΔNTE was crystallized in a monomeric form and the asymmetric unit contained three molecules of PK2ΔNTE.

The full-length PK2 forms a tetramer with a flower-like shape, with approximate dimensions of 90 Å × 100 Å × 60 Å (**Fig. 1b** & **1c**). The four molecules of the PK2 are extensively intertwined at the center of the complex through the four NTE regions of PK2 protomers. The NTE regions compose a globular connecting module with approximate dimensions of 30 Å x 25 Å x 20 Å. We named the connecting module as <u>NTE</u>-<u>tet</u>ramerization domain (hereafter NTE-TET domain, **Fig. 1c** & **Extended Data Fig. 1a**). The EKCM domains of the four PK2 molecules are located outside of the complex and surrounds the NTE-TET domain (**Fig. 1b** & **1c**). The complex comprises a 30 Å deep crevice at the center of the complex and its surface is negatively charged (**Extended Data Fig. 1b & 1c**). The *B*-factor values of the central region of the tetramer including the NTE-TET domain are approximately 30 Å^2^, comparable to the core of the EKCM domains of the protomers, and this low *B*-factor values of the NTE-TET domain indicates that the tetrameric organization is in a rigid state (**Extended Data Fig. 1d**). Therefore, the intertwinement of the NTE-TET domain is robustly associated and serves to constitute the core of the PK2 homotetramer.

### Tetrameric structure of PK2

The PK2 homotetramer is assumed to be composed of a dimer of two dimers, Mol-AB and Mol-A’B’, with a C2 symmetry (**Fig. 1b** &**Extended Data Fig. 1e**). Within the homotetramer, the PK2 protomers adopt two different conformations, Mol A and Mol B at non-equivalent positions, and Mol A’ and Mol B’ at their symmetrically related positions, respectively (**Fig. 1b** & **Extended Data Fig. 1e**). The PK2 protomers of Mol A and Mol A’ have a similar structure (root-mean-square deviation [rmsd] of 0.35 Å for 208 equivalent Cα atoms), and those of Mol B and Mol B’ have also similar structure (rmsd of 0.58 Å for 209 equivalent Cα atoms). Therefore, in this paper, we describe the structural features of Mol A and Mol B unless otherwise stated. The EKCM domains of the PK2 protomers, Mol A and Mol B, are in a dimeric form with a pseudo two-fold axis (**Extended Data Fig. 1e**). The interaction between the two EKCM domains of Mol A and Mol B is mediated by extensive binding, whereas that of Mol A and Mol B’ is only through one binding site (**Extended Data Fig. 2**). Although the two EKCM domains of Mol A and Mol B form the pseudo-symmetric dimer, the NTEs of Mol A and Mol B are clearly asymmetric because of their different conformations, as described below.

PK2 protomers have two distinct domains, the NTE region (residues 1-22) forming the NTE-TET domain and the C-terminal core region (residues 23-210) corresponding to the EKCM domain (**Fig. 1d**). Notably, a superposition of Mol A and Mol B based on the EKCM domains showed that the NTE regions of Mol A and Mol B have completely different conformations, despite having the same amino acid sequence (**Fig. 1d & Extended Data Fig. 1f**). The NTE region of Mol A contains one β-strand (β^MolA^), while the NTE region of Mol B instead contains one α-helix (α^MolB^) and one β-strand (β^MolB^). When Mol A and Mol B are superimposed, the distances between the corresponding Cα atoms of Mol A and Mol B in the NTE region are very far, and its N-terminus of Mol A is approximately 30 Å away from that of Mol B (**Extended Data Fig. 1g**). These alternative conformations adopted by the NTE enable PK2 to form the rigorously intertwined NTE-TET domain for the assembly PK2 homotetramer. Structural searches using the DALI server (33) could not find any structure homologous to the NTE-TET domain. Thus, the configuration of the NTE-TET domain appears to be the novel α/β fold specific to PK2 that contributes to its homotetrameric formation.

Contrary to the NTE, the EKCM domains of Mol A and Mol B retain similar structures (rmsd of 0.59 Å for 185 equivalent Cα atoms) comprised of one pair of antiparallel β strands (β1, β2), nine α helices (α1-α9) and three 3_10_ helices (η1-η3) (**Fig. 1d, Extended Data Fig. 3**). As reported previously, the amino acid sequence of the EKCM domain shares significant sequence homology with the C-lobe of eIF2α protein kinase domains (27, 29). The structural similarity search of the EKCM domain using the DALI server (33) showed that the most similar structures are found in the eIF2α kinases, including the GCN2 from *Saccharomyces cerevisiae* (PDB code: 1zxe, *Z*-score = 21.6, rmsd = 2.3 Å) and *Homo sapiens* (PDB code: 6n3o, *Z*-score = 20.1, rmsd = 2.3 Å), the PERK from *Mus musculus* (PDB code: 3qd2, *Z*-score = 21.6, rmsd = 2.0 Å) and *H. sapiens* (PDB code: 4x7k, *Z*-score = 19.1, rmsd = 1.7 Å), and the PKR from *H. sapiens* (PDB code: 2a1a, *Z*-score = 20.4, rmsd = 2.0 Å). Notably, residues 98-108 of PK2, corresponding to the activation segment of the eIF2α kinase domain (34, 35), contains one helix α3 and adopt a compactly folded conformation (**Extended Data Fig. 4a-4c**). Additionally, the loop between α1 and α2 (α1-α2 loop) show an extended conformation compared to the other kinases (**Extended Data Fig. 4d**). These two regions are directly involved in the tetramerization of PK2, and therefore these structural characteristics of PK2 may have been acquired to accommodate the homotetramer formation during the viral evolution.

The crystal structure of PK2ΔNTE, the isolated EKCM domain, showed that the structures of PK2ΔNTE in the asymmetric unit adopt basically similar conformations (rmsd of 0.41-0.42 Å for 187 equivalent Cα atoms), and are well-superimposable to the EKCM domain of the full-length PK2 (**Fig. 1d**). The globular structure of the PK2ΔNTE suggested that the NTE region is not required for the fold and maintenance of the structure of EKCM domain. Furthermore, the monomeric behavior of the truncated protein PK2ΔNTE indicates that the NTE of the PK2 protein has a crucial role for the homotetramerization of the full-length PK2, consistent with the analysis using the size exclusion chromatography (SEC) (32).

To analyze the homotetramer/monomer transition of PK2, we evaluated the full-length PK2 by size exclusion chromatography. As shown previously, the full-length PK2 was separated by SEC into two peaks, corresponding to the tetramer and monomer forms (32). After the 1^st^ SEC analysis of PK2, the peaks of tetramer and monomer were collected separately, and each was reapplied to the SEC. The SEC analyses showed that the individual peaks of PK2 were again separated into the two peaks, corresponding to the sizes of tetramer and monomer (**Fig. 1e**). Therefore, the SEC results indicated that the PK2 exists as reversible tetramer-monomer equilibrium in solution. The result also indicated that the ratios of the tetramer and monomer differed in the tetramer-applied and monomer-applied cases. The majority of the tetramer fraction of 1^st^ SEC remained in the tetramer (89%). However, the monomer fraction of 1^st^ SEC existed as a mixture of comparable amounts of tetramer (38%) and monomer (62%). Therefore, although the tetramer–monomer transition of the PK2 protein is a reversible transition, the tetrameric form of PK2 would be relatively more stable than the monomeric form of PK2. For the eIF2α kinase inhibition, the monomeric form of PK2 would preferentially bind to the kinase domain (32).

### Structural determination of PK2-BmHRI complex

To understand the molecular mechanism of eIF2α kinase inhibition by PK2, we next aimed to determine the complex structure of PK2 and eIF2α kinase by X-ray crystallographic analysis. First trials of the complex crystallization between PK2 and longer eIF2α kinase regions, including the kinase domain of PKR (PKR^KD^) and N-lobes of PKR and BmHRI, had failed to yield those crystals. Then, multiple trials of truncation, coexpression, copurification and crystallization screening experiments with BmHRI N-lobe and full-length PK2 were conducted. Crystals of co-purified complexes of PK2 and BmHRI were obtained with subdomain III-IV of BmHRI, which is the essential element for interaction with PK2 (**Fig. 2a**). The crystals of the complex contained the full-length PK2 and the subdomain III-IV of BmHRI, including residues 188 to 209 (form I) and residues 188 to 213 (form II), with an N-terminal MBP-tag to facilitate crystallization (**Fig. 2b & 2c**). The complex structures of PK2 and BmHRI, form I and form II, were refined to 2.0 Å and 2.1 Å resolutions with *R* factors of 19.7% (*R*_free_=22.4%) and 21.4% (*R*_free_=24.4%), respectively (**Extended Data Table 1 & Extended Data Fig. 5**). The two complex structures, form I and form II, exhibited basically similar complex formation and molecular interactions between PK2 and BmHRI. We therefore describe the structural features of the complex form II unless otherwise stated.

**Figure 2.**
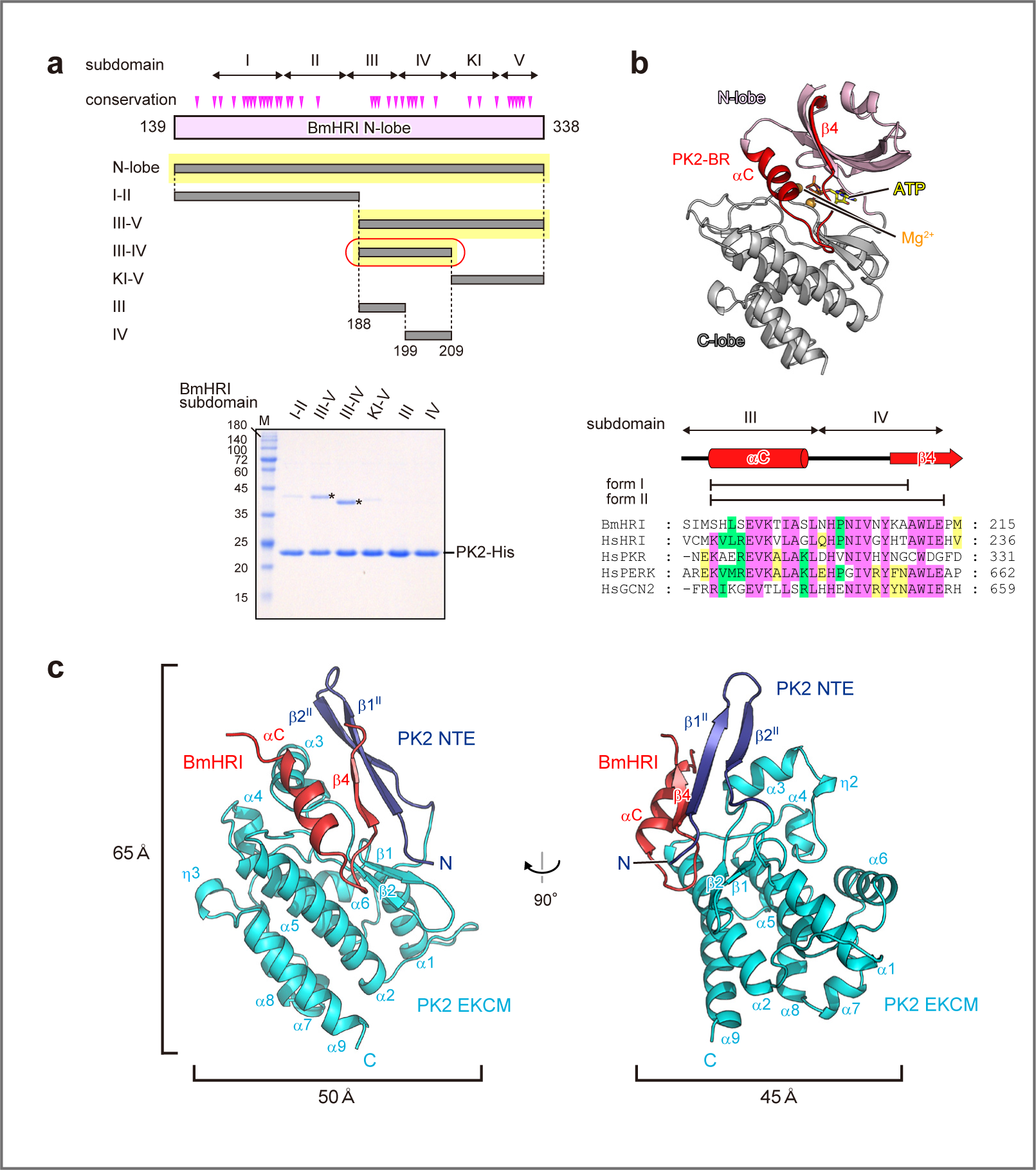
Complex structure of PK2 and BmHRI. **(a)** Schematic representation of BmHRI N-lobe showing the kinase subdomains on the top row, and the amino acid conservation between BmHRI and PKR in the next row. Identical amino acid residues between BmHRI and PKR are shown by pink arrowhead. The gray bars indicate the BmHRI variants used in this study. The yellow shadows show the variants bound to PK2, and the bar encircled by a red line shows the variant that yielded the complex crystal. The coexpression and purification of His-tagged PK2 and MBP fusion BmHRI variants is shown in the lower panel. The Ni-column purified samples were separated by SDS-PAGE and stained with Coomassie Brilliant Blue (CBB). Asterisks indicate the BmHRI variants bound to PK2. **(b)** Ribbon representation of human PKR^KD^ (PDB ID: 2A19) showing the region corresponding to PK2-BR, N-lobe and C-lobe, colored red, pink and gray, respectively. The amino acid sequence alignment of PK2-BR with eIF2α kinases is shown in the lower panel. Highly conserved residues are shaded in pink, while moderately and weakly conserved residues are shaded in green and yellow, respectively. **(c)** Ribbon representation of PK2-BmHRI complex. The NTE region and EKCM of PK2 are colored navy blue and cyan, respectively. The PK2-BR of BmHRI is colored red. Secondary structures are labeled as shown in Extended Data Fig. 3.

The complex structure reveals a globular complex of PK2 and BmHRI in a 1:1 stoichiometry with approximate dimensions of 65 Å × 50 Å × 45 Å (**Fig. 2c**). The PK2-binding region of BmHRI, named here as PK2-BR, consists of residues 188-213 and corresponds to the subdomain III-IV of eIF2α kinase, which contains αC helix and β4 strand. PK2-BR intrudes into a cleft between the NTE region and EKCM of PK2. The entire region of the PK2-BR of BmHRI, including a loop between αC helix and β4 strand (αC-β4 loop), makes extensive interactions with the PK2 cleft between the NTE region and EKCM. Notably, the NTE structure of the complex adopts the binding form different from the protomers, Mol A and Mol B, of the homotetramer of apo-PK2 (**Fig. 1d**). Instead, the NTE of the complex adopts a β-hairpin structure composed of two extended β-strands, β1^II^ and β2^II^, forming an antiparallel β-sheet with β4 of BmHRI (**Fig. 2c**). The PK2-BR is one of the characteristic regions containing residues that are conserved among the eIF2α kinase family (**Fig. 2b**, 35), consistent with the biological target selectivity of PK2 (27, 29), as discussed below.

### Complex formation between PK2 and BmHRI

The BmHRI PK2-BR, including αC helix and β4 strand, is localized in the groove between the NTE and EKCM domain (**Fig. 2c & 3a**), and the relative position of PK2-BR in the complex is similar to that of the subdomain III-IV in the eIF2α kinase PKR (compare **Fig. 3a & 3b**). The EKCM domain has similar conformation with those of the apo-PK2 with RMSD values lower than 0.66 Å, which suggests that the EKCM domain interacts with BmHRI without large conformational change. On the other hand, the NTE of the complex adopts an extended β-hairpin structure that is quite different from those within the homotetrameric structure of PK2. The NTE β-hairpin consisting of β1 (β1^II^) and β2 (β2^II^) of the form II complex comprises a continuous intermolecular β-sheet with the β4 of BmHRI in the order of β4^BmHRI^ -β1^II^ -β2^II^ (**Fig. 3a & 3c**). Intriguingly, the β-sheet orientation of the intermolecular interaction of β4^BmHRI^ is different from that of the intramolecular interaction of β4 within PKR^KD^. In the PKR^KD^, strands β4 and β5 form an antiparallel β-sheet; instead in the PK2-BmHRI complex, the strands β4^BmHRI^ and β1^II^ form a parallel β-sheet (**Fig. 3a-3d**). The β-hairpin consisting of β1^II^ and β2^II^ of PK2 NTE occupies a space roughly corresponding to β5 and β3 of N-lobe of PKR^KD^ (**Fig. 3a & 3b**). Therefore, the complex structure implies that the NTE, in cooperation with EKCM domain, captures and isolates the PK2-BR of BmHRI, leading to the unfolding of the kinase N-lobe as well as the inability of ATP-binding and the phosphorylation activities.

**Figure 3.**
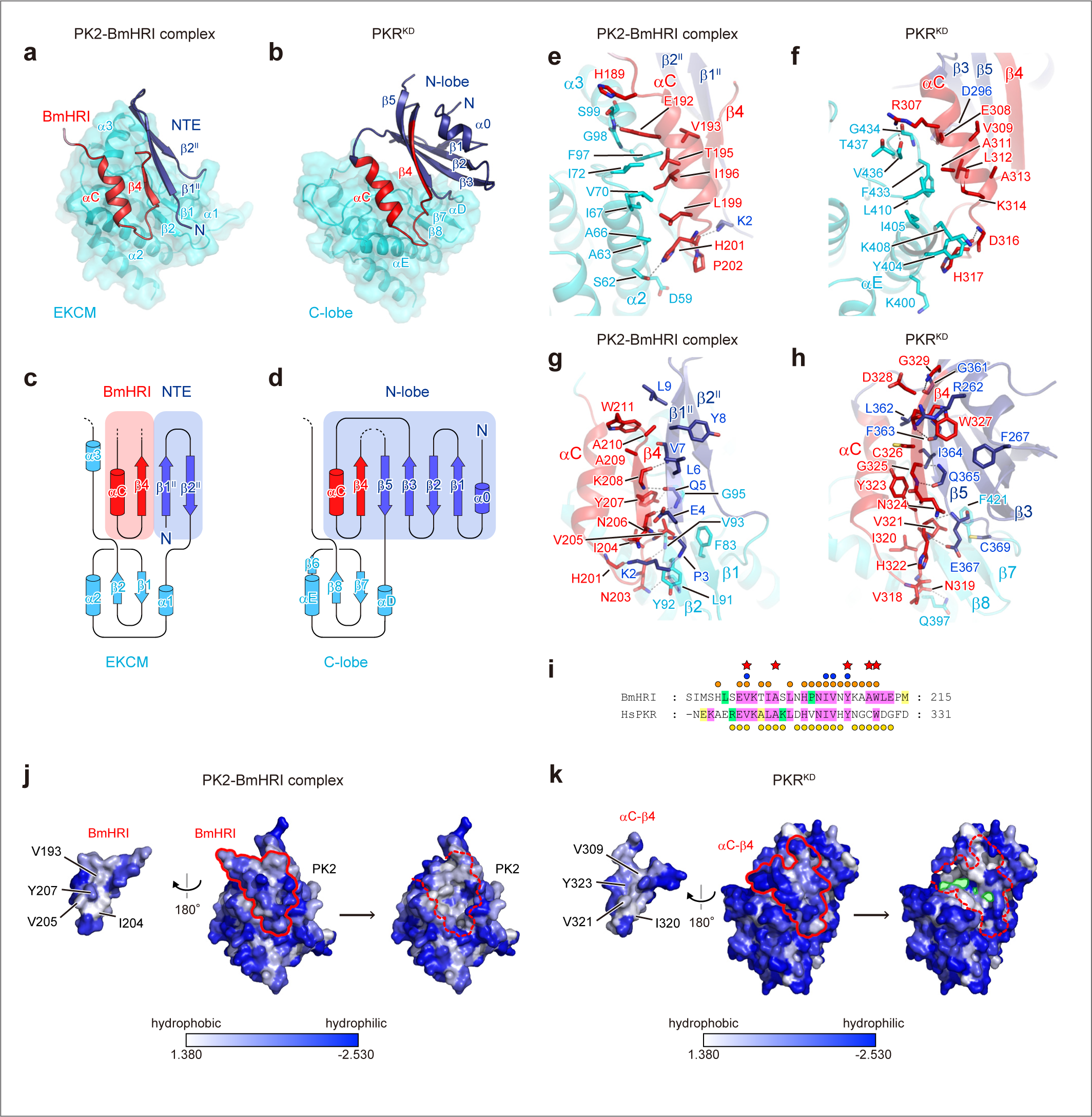
Contact interface between PK2 and BmHRI. **(a)** Ribbon representation of PK2-BmHRI complex. The NTE region and EKCM of PK2 are colored navy blue and cyan, respectively. The PK2-BR of BmHRI is colored red. A transparent surface of EKCM is shown. **(b)** Ribbon representation of PKR^KD^ (PDB ID: 2A19). The subdomain III-IV, corresponding to the PK2-BR, is colored red, while other parts of N-lobe and C-lobe are colored navy blue and cyan, respectively. A transparent surface of C-lobe is shown. **(c)** Topology diagram of the interface of PK2-BmHRI complex colored as in Fig. 3a. **(d)** Topology diagram of PKR^KD^ of the intramolecular interface of subdomain III-IV colored as in Fig. 3b. **(e)** A close up view of the interface between PK2 and BmHRI PK2-BR from the left side of Fig. 3a. Residues forming the interface are shown as stick model and colored as in Fig. 3a. **(f)** A close up view of the intramolecular interface between subdomain III-IV and the rest of PKR^KD^ from the left side of Fig. 3b. Residues forming the interface are shown as stick model and colored as in Fig. 3b. **(g)** A close up view of the interface between PK2 and BmHRI from the right side of Fig. 3a. **(h)** A close up view of the intramolecular interface between subdomain III-IV and the rest of PKR^KD^ from the right side of Fig. 3b. **(i)** A sequence alignment of the subdomain III-IV of BmHRI and HsPKR shaded as in Fig. 2b. Orange circles indicate the interface residues of the PK2-BmHRI complex, and yellow circles indicate the intramolecular interface of the subdomain III-IV and the rest of N-lobe. Red stars indicate the conserved residues amongst the eIF2α kinase family (35). Blue circles indicate the residues substituted in the mutation experiment. **(j)** Hydrophobic surface representation of the interface of the PK2-BmHRI complex colored according to the scale shown at the bottom. The surface of BmHRI PK2-BR and PK2 are shown on the left and right, respectively. **(k)** Hydrophobic surface representation of the intramolecular interface of the subdomain III-IV and the rest of PKR^KD^. The surface of the subdomain III-IV and the rest of PKR^KD^ are shown on the left and right, respectively.

The interactions between BmHRI PK2-BR and PK2 are mainly hydrophobic, except for the β-sheet between β4^BmHRI^ and β1^II^ (**Fig. 3e & 3g**). The αC helix of PK2-BmHRI complex is located on the upper space between the α2 and α3 of EKCM domain (**Fig. 3e**). The conserved histidine residue His201 of BmHRI forms a hydrogen bond with the side chain of Ser62 of EKCM domain, and the main chain oxygen of His201 also forms a hydrogen bond with the side chain of Lys2 of NTE region. The hydrophobic interactions were found throughout the αC helix of BmHRI, which is similar to the intramolecular interactions of αC helix within the PKR^KD^, though α3 helix is absent in PKR^KD^ (**Fig. 3e & 3f**). In the interactions of β4^BmHRI^, the side chains of Asn203 and Tyr207 formed hydrogen bonds with the main chains of Leu91 and Gly95, respectively (**Fig. 3g**). The electron densities of Leu212 and Glu213 of BmHRI were missing, and the side chain of Trp211 was flipped to α3 helix compared to the corresponding residue Trp327 of PKR^KD^ (**Fig. 3g & 3h**). The flipping of Trp211 may induce the structural change of the C-terminal end of β4^BmHRI^, which causes the shortening of strand moiety of β4^BmHRI^ in the PK2-BmHRI complex.

In the complex structure of PK2 and BmHRI, the residues involved in the complex interface spanned the entire subdomain III-IV, and importantly, most of the residues conserved specifically in the eIF2α kinase family participated in the interface of the complex (**Fig. 3i**, 35). This commonality is well compatible with the biological function of PK2. To compare the intermolecular interaction of PK2-BmHRI complex with the intramolecular interaction in PKR^KD^, the molecular surfaces of the interfaces were colored according to hydrophobicity (**Fig. 3j & 3k**). These showed that both interactions are mainly hydrophobic interactions. In the complex formation of PK2 and BmHRI, the intermolecular surface is completely closed (**Fig. 3j)**. However, the intramolecular interface of αC helix-β4 strand in PKR^KD^ has cavities for ATP binding (**Fig. 3k**), which exhibit hydrophilic property both in the ATP-bound state and ATP free form. Therefore, the complex formation between BmHRI PK2-BR and PK2 may be more stable than the intramolecular interactions between the subdomain III-IV and the rest of kinase domain, which may lead to the favorable complex formation for the kinase inhibition.

### Biochemical characterizations of PK2 and eIF2α kinase complex

To test whether the PK2 and eIF2α kinase complex exhibits the kinase activity, we employed in vitro biochemical assay using the pre-bound eIF2α kinase PKR^KD^-PK2 complex and PKR^KD^ alone. The result showed that PKR^KD^ exhibits the kinase activity with the Km value of 429.7 μM (**Fig. 4a**), and the value is different from the previously reported Km (150 μM, 23), which may be due to the differences in the assay conditions. On the other hand, the PKR^KD^-PK2 complex exhibited no detectable kinase activity (**Fig. 4a**). This indicates that the binding of PK2 to PKR^KD^ leads to a complete loss of its kinase activity.

**Figure 4.**
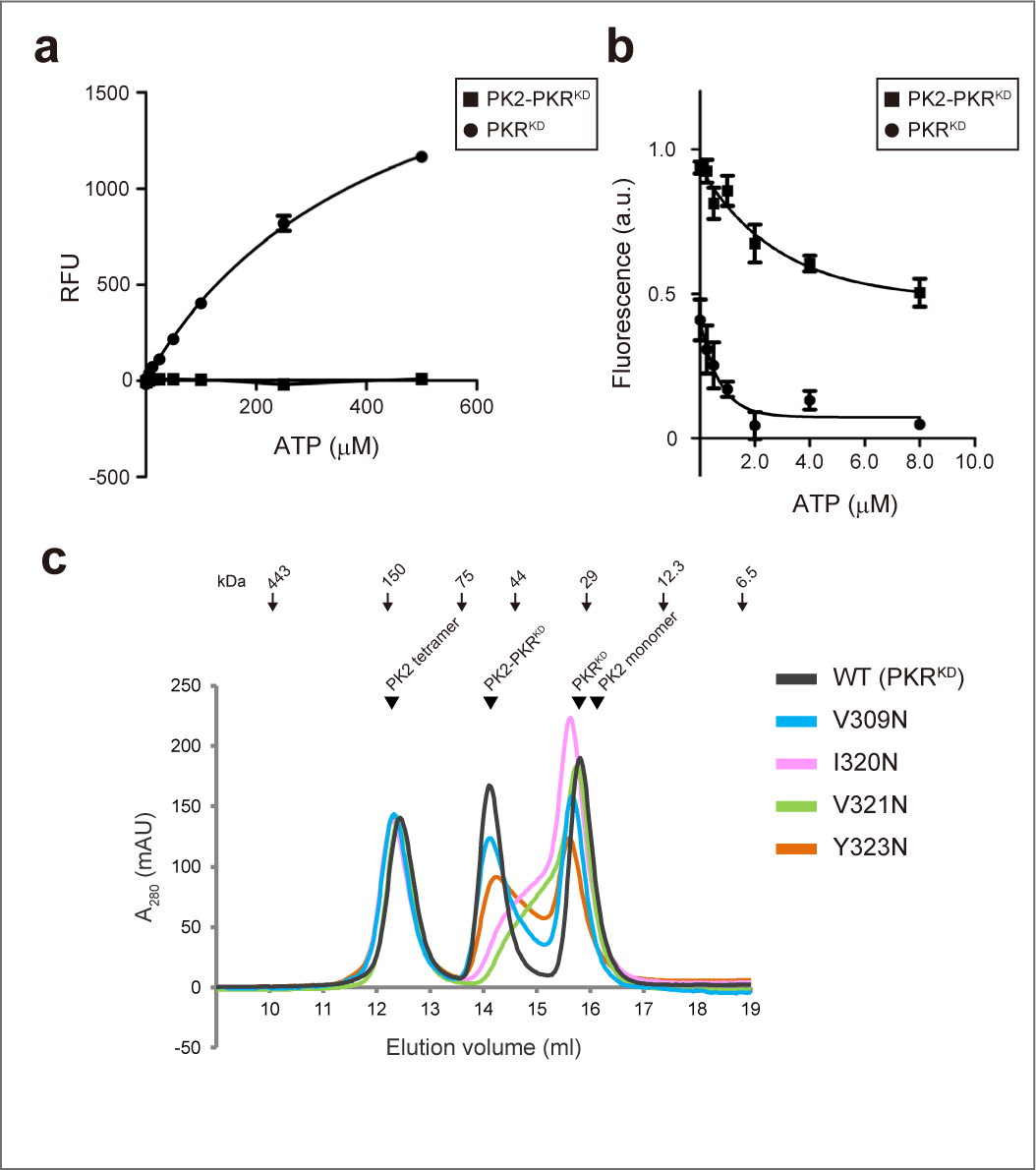
Biochemical characteristics of PK2-eIF2α kinase complex and its formation. **(a)** PKR^KD^ and PK2-PKR^KD^ complex were incubated with eIF2α, and ATP concentration was varied as indicated. Generated ADP through the kinase reaction was quantified as fluorescent signals. The assay was repeated three times. Error bars indicate the standard deviations. **(b)** Competition between fluorescent TNP-ATP and ATP for binding to PKR^KD^ and PK2-PKR^KD^ complex. ATP concentration was varied as indicated. The assay was repeated three times. Error bars indicate the standard deviations. **(c)** Representative SEC analysis of mixtures of PK2-PKR^KD^ complex, and PK2-PKR^KD^ mutant complexes as shown in legend.

Next, we examined the ATP binding affinities of PKR^KD^-PK2 complex and PKR^KD^ alone using TNP-ATP fluorescence and ATP titration (**Fig. 4b**, 36, 37, 38). The fluorescence value of TNP-ATP bound to PKR^KD^-PK2 complex was higher than that of TNP-ATP bound to PKR^KD^ (values at 0 m*M* ATP in **Fig. 4b**), which may be due to changes in the environment around the TNP-ATP binding site caused by the complex formation. The data of PKR^KD^ alone indicated the competition of ATP for TNP-ATP bound to PKR^KD^ (Ki^ATP^ 0.997 μM). However, the data of PKR^KD^-PK2 complex showed a weak competitive activity of ATP for TNP-ATP, and the fluorescence values remained at the high ATP concentration (**Fig. 4b**). The changes of the TNP-ATP fluorescence due to the complex formation may indicate that the PKR^KD^-PK2 complex lost its correct ATP binding activity by altering the ATP binding site. Consistent with this assay, previous biochemical study had shown that wild-type PK2 inhibits BODIPY-γ-S ATP displacement activity of PKR^KD^ with an IC_50_ of 30 μM (29). Taken together, these data suggests that the complex formation of PKR^KD^ with PK2 altered the structure of the ATP-binding site, thereby attenuating its ATP-binding activity. This is well compatible with our structural observation showing the extensive interaction between PK2 and the BmHRI subdomain III-IV, that includes a crucial regulatory element for ATP binding in kinase catalytic reaction.

To analyze the PK2 complex formation of wild-type PKR^KD^ and its mutants, SEC analysis was performed as in the previous complex analysis (**Fig. 4c**, **Extended Data Fig. 6,** 32). In the experiment, the hydrophobic residues of PKR^KD^ on the presumptive surface of the complex were substituted to hydrophilic residue asparagine. The residues, V309, I320, V321 and Y323 of PKR^KD^ corresponding to V193, I204, V205 and Y207 of BmHRI, respectively, would be situated on the intermolecular surface bound to PK2 (**Fig. 3i-3k**). Only monomeric form of PK2 can form complex with PKR^KD^ (32), and thus the peak of PK2 tetramer is not affected by incubation with PKR^KD^ (**Fig. 4c**). The results of the PK2 complex formation with WT PKR^KD^ showed that the chromatogram exhibited three clearly separated peaks corresponding to the PK2 tetramer, the PK2-PKR^KD^ complex, and the PKR^KD^ monomer (**Fig. 4c**). However, the chromatogram of the PK2 mixed with mutant proteins, V309N and Y323N, reduced the peak of the complex of PK2 and PKR^KD^, and these peaks decreased slowly compared to that of the WT PKR^KD^. Furthermore, in the case of I320N and V321N, the PK2-complex peaks were significantly lower, indicating low efficiency of the complex formation. The decrease in the peak of the complex and the peak tailings of PK2 mixed with PKR^KD^ mutants would indicate unstable complex formation between PK2 and PKR^KD^ mutants. The results of the SEC analysis of PKR^KD^ mutants are consistent with the result of the complex structure analysis.

### Conformational changes for PK2-BmHRI complex formation

Our crystallographic analysis revealed that PK2 adopts dramatically altered conformations between the homotetrameric forms and its complex form with BmHRI. These conformations of PK2 were compared by the superposition based on the EKCM domain (**Fig. 5a**). The conformational changes were mainly found in the NTE region. The NTE region of the protomer Mol A of PK2 tetramer contains one β strand, while that of Mol B contains one α helix and one β strand. In the BmHRI complex, the NTE region contains a β hairpin consisted of two β strands. The spatial arrangement of each NTE is also different, whereas the Cα atoms of Asn23 at the N-terminus of EKCM domain were essentially superimposable within these conformations (**Fig. 5a**). Therefore, these variable conformations of the NTE region would serve a dual function of the homotetramerization and BmHRI inhibition.

**Figure 5.**
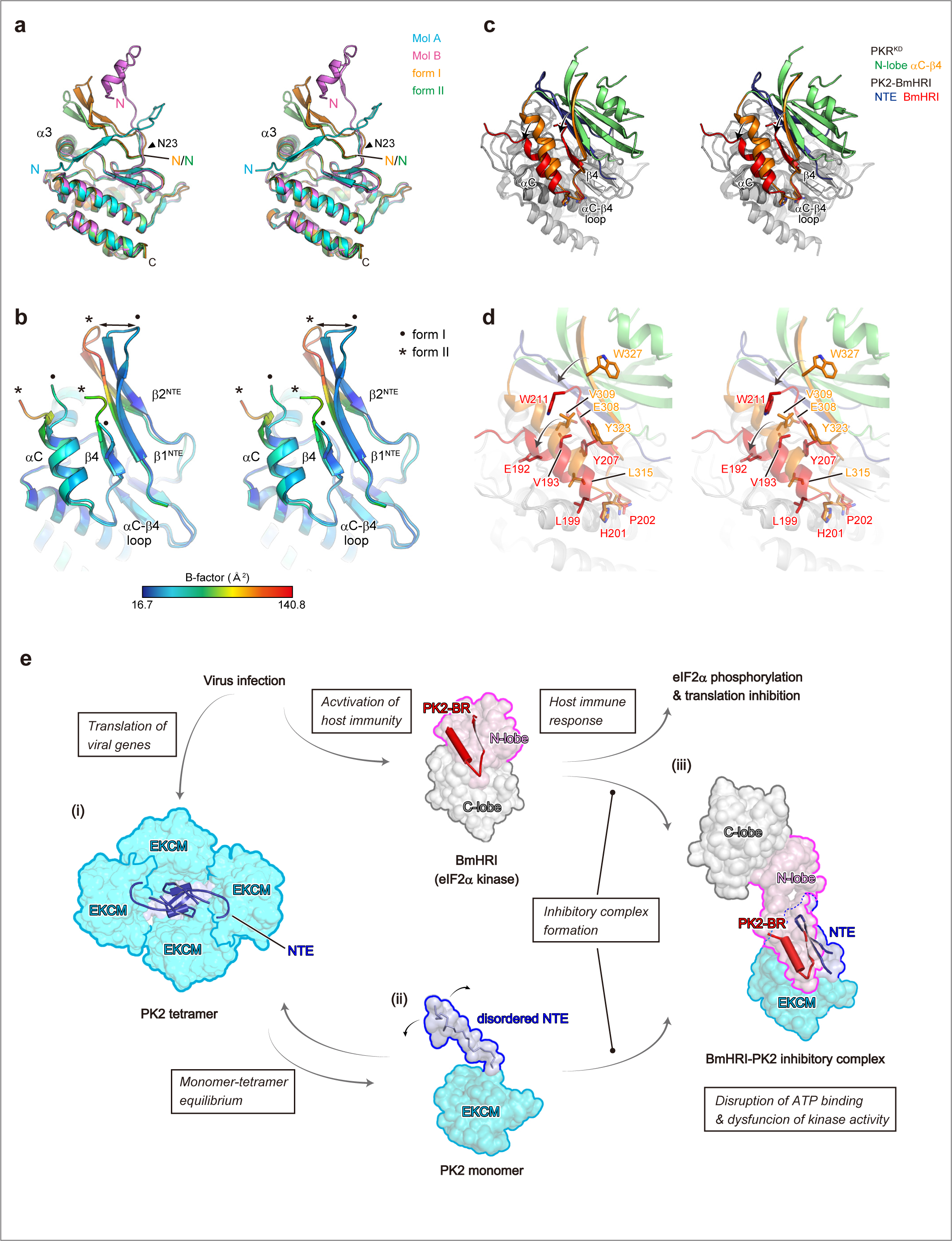
Conformation changes for the PK2-BmHRI complex formation and functional model for BmHRI inhibition by PK2. **(a)** Stereoview of superposition of PK2 protomers and PK2 complexed with BmHRI. Structures are colored as indicated in panel. **(b)** Stereoview of superposition of PK2 complexed with BmHRI, form I and form II, colored according to B-factor as shown in the spectrum bar. Structures of form I and form II are labeled as in panel. **(c)** Stereoview of superposition of PK2-BmHRI complex and PKR^KD^ based on the C-lobe/EKCM domain. Individual portions of the structures are colored as in panel. **(d)** Close-up stereoview of the superposition of PK2-BmHRI complex and PKR^KD^ around the subdomain III-IV. **(e)** Functional model of PK2 for BmHRI inhibition. PK2 encoded in viral genome is translated in the infected cell, and forms both homotetramer and monomer (i, ii). The functional monomer of PK2 dissociates from the homotetramer by thermodynamic driving forces. The PK2 monomer binds to PK2-BR of BmHRI and sequesters the kinase activity of the innate immune signalling (iii). The structure of BmHRI kinase domain was predicted by Alpha Fold2. The structure of disordered NTE of the PK2 monomer (ii) and the kinase domain of BmHRI except for the PK2-BR (iii) were modelled manually. The secondary structures of NTE and PK2-BR is illustrated in the ribbon model.

The conformations of PK2-BR and the NTE region in the complex form I and form II were compared by the superposition based on the EKCM domain and colored by B-factor values (**Fig. 5b**). The result showed that the upper half of the β hairpin of NTE is variable, and the Cα atoms of Arg12 of forms I and II are 9.6 Å apart from each other. Furthermore, the upper side of the αC helix and β4 strand showed higher β-factors in the complex form II. Therefore, the complex formation between PK2 and BmHRI would be achieved mainly by the interaction within the internal segments of αC helix and β4 strand, and β1^NTE^ and β2^NTE^, within the complex.

To analyze the conformational change of PK2-BR of BmHRI, the structures of PKR^KD^ and PK2-BmHRI complex were superimposed based on the C-lobe/EKCM domain (**Fig. 5c & 5d**). Overall, in the complex structure of PK2-BmHRI, PK2-BR had shortened toward the tip of the αC-β4 loop (**Fig. 5c & 5d**). In the complex structure of PK2-BmHRI, an additional helix was wound at the C-terminus of the αC helix. Leu199 of BmHRI was incorporated to the αC helix, but the corresponding residue L315 of PKR^KD^ was not (**Fig. 5d**). Therefore, the extension of the αC helix of the PKR^KD^ and BmHRI in complex with PK2 would be changeable. In the β4 strand, the side chains of Tyr207 and Trp211 of BmHRI were flipped toward the EKCM domain, compared to the corresponding residues Tyr323 and Trp327 of PKR^KD^. The distances of the side chains between Tyr207 of the BmHRI and Tyr323 of PKR^KD^ and between Trp211 of the BmHRI and Trp327 of PKR^KD^ are 18.0 Å and 12.7 Å, respectively. These conformational changes of both PK2 NTE and PK2-BR of BmHRI would allow the formation of the stable inhibitory complex.

## Discussion

### Functional model for inhibition of host eIF2α kinase immune signalling by PK2

Here, we present the X-ray crystal structures of PK2 both in the apo form and in the complex with BmHRI at resolutions of 2.0-2.8 Å. The series of structural and biochemical analyses provide mechanistic insights into the inhibition of host eIF2α kinase BmHRI by PK2, which explain how virus adapts to host cell for proliferation. Based on the data, we propose a functional model of PK2 to inhibit BmHRI immune signalling as follows (**Fig. 5e**). Upon viral infection, PK2 encoded in viral genome is transcribed and translated in the infected cells. PK2 exists in both homotetrameric and monomeric forms, with reversible transition between the homotetramer and monomer. In the homotetramer, the NTE region adopts two conformations of Mol A and Mol B, both of which are involved in homotetramerization (**Fig. 1b-1d**). The function of the PK2 homotetramerization would be to provide stability to its NTE structure, which is a structurally changeable region and required for inhibitory complex formation with BmHRI (**Fig. 2c**). Generally, oligomerization of protein increases its stability, and the reduction of surface area of the monomer can protect it from denaturation (39, 40, 41), whereas a disordered region decreases protein half-life in living cells (42, 43, 44). Therefore, the PK2 homotetramerization and the NTE sequestration would increase the stability of PK2, and functional monomer of PK2 would dissociate from the homotetramer by thermodynamic driving forces, as observed in SEC analysis (**Fig. 1e**). Then, the monomeric form of PK2 specifically binds to host eIF2α kinase BmHRI, generating the inhibitory complex with a 1:1 molecular ratio (**Fig. 2c**). The complex formation of BmHRI with PK2 alters the ATP-binding site and abrogates its kinase activity, leading to shut-off of the host immune signalling (**Fig. 4a & 4b**).

The mutation analyses presented here are consistent with the complex structure of PK2-BmHRI, and the previous mutation analyses are also compatible with the molecular interaction of the inhibitory complex. In the previous analyses, PK2 mutations, F18A (since it is a variant, Phe18 is Ile18 in this study) and Δ1-21 (ΔNTE), impaired PKR binding and its inhibitory function (29). In the complex structure, Ile18 positions at the middle of the β2^NTE^ and makes hydrophobic interaction with Val7 of the β1^NTE^ that lies at the complex interface. Therefore, the impairments of PK2 F18A would be an indirect effect for the inhibitory complex formation. Taken together, the structural data of the complex between PK2 and BmHRI is well aligned with the results of these biochemical assays.

### PK2 binds to subdomain III-IV for eIF2α kinase inhibition

Viral PK2 directly binds to and inhibits the kinase domain of eIF2α kinase, thereby stimulating the viral replication in the infected cells (27, 28, 29). The truncation experiments of eIF2α kinase BmHRI revealed that PK2-BR, the subdomain III-IV including αC helix and β4 strand, is the indispensable region for PK2 binding (**Fig. 2a**). Taken together, the viral strategy of targeting PK2-BR to inhibit the host immune signalling of eIF2α kinase seems to be rational for the following reasons. First, the PK2-BR contains invariant residues that are strictly conserved within the eIF2α kinase family (35), and these residues are indeed involved in the interaction between PK2 and BmHRI (**Fig. 3**). The complex structure between PK2 and BmHRI PK2-BR showed that these conserved residues of eIF2α kinase, Val193, Tyr207, Ala210, and Trp211 of BmHRI, form the interface of the complex (**Fig. 3e & 3g**). For the selective inhibition of the host eIF2α kinase, viral PK2 must specifically bind to and inhibit the eIF2α kinase, not other kinases in the cell (27, 29). The PK2-BR contains the residues specifically conserved among eIF2α kinases (**Fig. 3i**), consistent with the molecular function of PK2 as a viral proteinaceous inhibitor of the host eIF2α kinase. Second, the subdomain III-IV contains the αC helix, which is an important regulatory element of conformational changes for its catalytic activity (45, 46, 47, 48). The αC helix, the only conserved helix in the β-strand rich kinase N-lobe, includes the strictly conserved glutamate residue (Glu91 of PKA, Glu192 of BmHRI), that forms an ion pair with the lysine (Lys72 of PKA, Lys174 of BmHRI), which coordinates the α and β phosphates of ATP. The αC helix also interacts with the N-terminal region of the activation loop, and its conformation affects the flipping of the conserved Asp-Phe-Gly (DFG) motif. These interactions link the conformation of the helix to the ATP binding for its kinase activity (46, 48). In addition, αC helix participates in the highly conserved and important intramolecular network, regulatory spine (R-spine), mainly composed of hydrophobic residues. The movement of αC helix disrupts the R-spine hydrophobic network, resulting in the kinase inactivation (45, 47, 49). Moreover, the αC helix and β4 strand in the eIF2α kinases, PKR^KD^ and GCN2^KD^ form the dimer interface, which promotes its catalytic activity (34, 35). Therefore, the mechanism of the sequestration and blockade of the BmHRI subdomain III-IV, including αC helix and β4 strand, by PK2 illustrates the physiological function of suppression of the host eIF2α-kinase immune signalling.

### Evolution of viral PK2 through horizontal gene transfer from host genome

The analysis of the amino acid sequence of PK2 indicated that the PK2 showed the highest sequence similarity to the C-lobe of eIF2α kinase domains, and the top hits corresponded to the HRI orthologs of insects (29). In addition, the phylogenetic analysis of PK2 genes and eIF2α kinases demonstrated that PK2 closely associates with the insect HRI orthologs, supporting a model of horizontal gene transfer from an insect to a viral genome, in which the kinase has been partially evolved to inhibit the host immune system (29). Based on this model and the complex structure presented here, we could evaluate the adaptation and evolution of PK2 to inhibit host BmHRI kinase. The multiple sequence alignment of PK2 proteins and eIF2α kinase domains displayed the homology between the PK2 NTE regions and eIF2α kinase subdomain I (**Extended Data. Fig. 3**). This homologous region of the PK2 NTE and the eIF2α kinase subdomain I, both showing a structural moiety of a β-hairpin, may suggest the origin of the NTE region. Given the homology between the PK2 NTE and the kinase subdomain I, the full-length PK2 might has emerged by a deletion of the internal sequence between the subdomain II and the kinase insert after the horizontal gene transfer of an HRI ortholog. The β-hairpin of NTE directly contacts with β4 of BmHRI, whereas the β-hairpin of the subdomain I indirectly interacts with β4 within the kinase N-lobe (**Fig. 3a-3d**). Therefore, after the deletion of the internal sequence, the NTE of PK2 might have evolved to selectively recognize host BmHRI and inhibit its immune signalling under positive selection pressure for the viral adaptation.

In the PK2 EKCM, the residues interacting with BmHRI are located on the α2 helix and around the β2 strand, which correspond to αE helix and β8 strand of eIF2α kinase, respectively (**Fig. 3e-3h**). The magnesium-binding DFG motif of the kinase was converted to the MFG motif, including Phe97 and Gly98, in the PK2 sequence (29, **Fig. 3e**), which comprise of the complex interface. The MFG motif lies on the top of the α3 helix that is unique to the PK2 structure (**Fig. 3e**). Thus, the α3 helix may be acquired for the formation of the PK2-BmHRI complex. Compared to insect HRI (BmHRI) C-lobe, which is considered to be the closest ancestral protein of PK2, six of the 16 residues within the EKCM domain that interact with BmHRI are conserved as identical, while the other residues are diverged (**Extended Data. Fig. 3**). Therefore, not only the PK2 NTE region but also the PK2 EKCM would have evolved to adjust and optimize the interface of the PK2-BmHRI complex.

### Viral inhibitors against host eIF2α kinase

As viral eIF2α kinase inhibitor, the vaccinia virus K3L has been well studied to inhibit human PKR (23–25). The vaccinia virus K3L is an eIF2α homolog that behaves as a pseudo-substrate and acts as a competitive inhibitor of PKR. The structures of apo form of K3L and PKR^KD^-eIF2α complex indicated that K3L and eIF2α share a homologous eIF2α/K3L fold, and K3L is therefore likely to interact with PKR^KD^ in a similar binding mode of the substrate eIF2α (23, 35). The complex model of K3L and PKR^KD^ implied that divergent structures in K3L, including the longer helix 1, helix 1-helix 2 connecting loop, helix 2, and the β1-β2 linker, confers PKR inhibitor function on K3L (35). Virus-associated RNA I (VA-I), encoded in adenovirus, competes with dsRNA to bind to PKR, and inhibits PKR autophosphorylation and activation (50–52). The crystal structure and biochemical analyses of VA-I RNA revealed that the overall structure of VA-I, essential for PKR inhibition, adopts an acutely bent shape and a central domain containing pseudoknot that may directly interact with the kinase domain for suppression of PKR activity (53). These structural analyses of K3L and VA-I implicated the functional segments and conformational change for inhibition of host eIF2α kinase. However, the detailed mechanisms of host eIF2α kinase inhibition via the interaction of the catalytic domain with viral inhibitors had remained enigmatic due to the absence of complex structure between the kinase domain and viral factor.

Here, we report for the first time the complex structure between host eIF2α kinase catalytic domain and viral inhibitor, revealing the novel molecular mechanism for the complexation leading to the kinase inhibition. The structural and biochemical analyses showed that viral PK2 binds specifically to the eIF2α kinase subdomain III-IV, the regulatory element for its kinase activity, as well as the conserved region among the eIF2α kinase family, which is well aligned with the functionality of the viral inhibitor. The sequestration of the subdomain III-IV, including αC helix and β4 strand, would inactivate the catalytic activity by disrupting of the β sheet in the kinase N-lobe. The newly formed continuous β-sheet between viral PK2 and host BmHRI is reminiscent of the complexation of MDA5 and virus V protein (54), in which the β hairpin structure of V protein disrupts the ATPase domain of MDA5. Similar to this viral inhibition mechanism, viral PK2 would unfold the eIF2α kinase N-lobe, resulting in the loss of the immune signalling activity. The structural analyses of viral PK2 and host eIF2α kinase advance our knowledge of the molecular interplay between the host immunity and viral countermeasure and elucidate the sophisticated mechanisms underlying the selective kinase inhibition by pseudo-kinase.

## Methods

### Protein expression, purification, and crystallization of PK2 and PK2ΔNTE

The PK2 from the baculovirus AcMNPV was expressed and purified as previously described (32). In brief, the stable mutant PK2 C181S/C211S with C-terminal His-tag was expressed at 25°C in *E. coli* cells and purified by Ni-NTA agarose column (Qiagen), HiTrap Q (Cytiva), and HiLoad 26/600 Superdex 200 column (Cytiva). The crystals of PK2 and SeMet-substituted PK2 for phase determination were obtained by sitting-drop vapour diffusion method (32). PK2ΔNTE was expressed and purified in a similar way to the full-length PK2 protein, and the crystal of PK2ΔNTE was obtained by a sitting-drop vapour-diffusion method (32). For X-ray data collection, the crystals were cryo-protected with the reservoir solution supplemented with 20% glycerol.

### Coexpression of PK2 and N-lobe regions of eIF2α kinase domain

The gene encoding PK2 C181S/C211S was cloned into pACYC Duet-1 to express the protein with C-terminal His-tag. The genes encoding N-lobe regions of eIF2α kinase domain of BmHRI were cloned into modified pMAL-c2x (55). The two plasmids, pACYC-PK2 and pMAL-N-lobe regions, were introduced into competent cells of *E. coli* BL21(DE3). The BL21(DE3) cells were grown at 37°C in LB medium containing 10 μg ml^-1^ chloramphenicol and 50 μg ml^-1^ ampicillin until the OD595 approximately 0.8, and then 0.2 m*M* IPTG was added to the medium and further cultured for 15h at 25°C. The cells were disrupted in lysis buffer [50 m*M* sodium phosphate pH 8.0, 300 m*M* NaCl, 5.0 m*M* β-mercaptoethanol (β-ME) and 10 m*M* imidazole] by sonication, and then centrifuged at 40,000g for 60 min. The supernatants were loaded onto Ni-NTA agarose column (Qiagen). The column was washed with a buffer containing 50 m*M* sodium phosphate pH 8.0, 300 m*M* NaCl, 5.0 m*M* β-ME and 20 m*M* imidazole, and the elution was carried out with the above buffer with 200 m*M* imidazole, and the elution fraction was analyzed by SDS-PAGE and CBB staining.

### Purification and crystallization of PK2 and BmHRI complex

The PK2 and BmHRI PK2-BR was coexpressed in BL21(DE3) cells. The sonication and purification with Ni-NTA agarose column (Qiagen) were conducted in a similar way to PK2 apo (32). Next, the Ni-NTA elution fractions were loaded on an amylose resin high flow column (New England Biolabs.), and elution was conducted with the same buffer with 10 m*M* maltose. The PK2 complexed with BmHRI PK2-BRwas further purified by Resource-Q chromatography (Cytiva), using 100 m*M*-1.0 *M* NaCl gradient. Finally, the complex was purified by HiLoad 26/600 Superdex 200 column (Cytiva) equilibrated with 20 m*M* Tris pH 8.0, 200 m*M* NaCl, 5 m*M* β-ME. The complex was concentrated using Amicon Ultra-15 centrifugal filter devices (Millipore) to 40 mg ml^-1^. A linker of two alanine residues between MBP and PK2-BR improved the quality of the complex crystal. The PK2 complexed with BmHRI residues 188 to 209 (form I) was crystallized using a reservoir solution of 1.8 *M* tri-ammonium citrate pH7.0, and the PK2 complexed with BmHRI residues 188 to 213 (form II) was crystallized using a reservoir solution of 0.2 M ammonium sulfate, 0.1 M HEPES pH 7.5, and 25% w/v polyethylene glycol 3,350 by a sitting-drop vapour-diffusion method. For X-ray data collection, the crystals were cryo-protected with the reservoir solution supplemented with 20% glycerol.

### Structural determination of PK2, PK2ΔNTE, PK2 and BmHRI complex

Single-wavelength anomalous dispersion (SAD) data were collected from the SeMet-labeled crystal of PK2. The phase of PK2 crystal was determined by SAD phasing at 2.8 Å and partial model was built by AutoSol (56) in PHENIX (57). An initial model of PK2 was built using the program Buccaneer (58). The initial model was iteratively refined using the program phenix.refine (59) and manually rebuilt using Coot (60) with the native data. The model contains chain A residues 1-209, chain B residues 1-209, chain C residues 1-212 and chain D residues 1-209. The structure was refined to R_factor_/R_free_ of 23.7%/29.9% to 2.7 Å resolution.

The phase of the PK2ΔNTE crystal was solved by molecular replacement with with Phaser (61) using the corresponding region of the full-length PK2 as the search model. The model was iteratively refined using the program phenix.refine (59) and rebuilt using Coot (60). The model contains chain A residues 23-210, chain B residues 23-212 chain C residues 23-209. The structure was refined to R_factor_/R_free_ of 22.7%/28.2% to 2.8 Å resolution.

The PK2-BmHRI complex crystals, form I and form II, were solved by molecular replacement with Phaser (61) from CCP4 suit (62) using MBP and PK2 EKCM domain as the search model. The crystal form I contains two complexes of PK2-BmHRI in the asymmetric unit cell, and the model contains MBP-fusion BmHRI (chains A and B) residues 1-392 (full-length MBP and BmHRI 188 to 209) and PK2 (chains C and D) residues 1-210. The crystal form II contains one complex of PK2-BmHRI in the asymmetric unit cell, and the model contains MBP-fusion BmHRI (chains A) residues 3-394 (full-length MBP and BmHRI 188 to 211) and PK2 (chain C) residues 1-210. The structures, form I and form II, were refined to R_factor_/R_free_ of 19.7%/22.4% to 2.0 Å resolution, and R_factor_/R_free_ of 21.4%/24.4% to 2.1 Å resolution, respectively.

The geometries of the final models of the full-length PK2, PK2ΔNTE and PK2-BmHRI complexes were evaluated using the program Molprobity (63). The data collection and refinement statistics of the PK2, PK2ΔNTE and PK2-BmHRI complexes are summarized in Supplementary Table 1.

### SEC analysis of PK2 and PKR kinase domain (PKR_KD_) complex

Size exclusion chromatography (SEC) analysis was performed using a Superdex 200 increase 10/300 column (Cytiva) with a buffer consisting of 20 m*M* Tris pH 8.0, 200 m*M* NaCl, and 5 m*M* β-ME at a flow rate of 0.4 ml/min with monitoring at 280 nm, in the same way as previously reported (32). Alcohol dehydrogenase, apoferritin (SIGMA) and gel filtration protein standards (Cytiva) were used for the calibration of the column. Full-length PK2 was applied to the Superdex 200 increase 10/300 column, and the elution fractions corresponding to the PK2 tetramer and monomer were collected separately. The two pools were concentrated separately using the Amicon Ultra-4 (Millipore), and re-applied to the Superdex 200 increase 10/300 column (Cytiva).

The kinase domain of an inactive PKR, PKR^KD^ ^K296R/C551A^, containing K296R/C551A mutations and β4-β5 loop deletion was prepared in a similar way in a previous study (32). The mixtures of PK2 and PKR^KD^ ^K296R/C551A^ or additional mutants were incubated on ice for 15 min, and were applied to the Superdex 200 increase 10/300 column (Cytiva) at a flow rate of 0.4 ml/min with monitoring at 280 nm.

### Preparation of PKR_KD_ and eIF2α proteins, and kinase activity assay

The kinase domain of an active PKR, PKR^KD^ ^H412N/C551A^, containing H412N/C551A mutations and β4-β5 loop deletion (35), was expressed as C-terminal His-tagged protein in the pET-22b vector. The expression and purification of PKR^KD^ ^H412N/C551A^ were conducted in a similar way to that of PKR^KD^ K296R/C551A mutant. For purification of PKR^KD^ ^H412N/C551A^, the anion-exchange chromatography using a Hitrap Q (Cytiva) was used instead of TOYOPEARL SuperQ-650. The complex of PK2 and PKR^KD^ was purified by a gel-filtration column, HiLoad 16/600 Superdex 200 (Cytiva) to remove excess PK2 and PKR^KD^. The *Saccharomyces cerevisiae* eIF2α (residues 3-175) was expressed as C-terminal His-tagged protein in the pET-26b vector. The expression and purification of eIF2α were conducted in a similar way to that of PKR^KD^ ^H412N/C551A^. For purification of eIF2α, HiLoad 16/600 Superdex 75 (Cytiva) was used instead of HiLoad 16/600 Superdex 200 (Cytiva).

The kinase assay was performed by measuring ADP production using the Fluorospark Kinase/ADP Multi-Assay Kit (Fujifilm Wako) on 384-well black plates (781900, Greiner Bio-One). PKR^KD^ ^H412N/C551A^ or PK2-PKR^KD^ ^H412N/C551A^ complex (1.0 µ*M*) were mixed with 0-500 µ*M* ATP and 10 µ*M* eIF2α in a buffer consisting of 20 m*M* Tris pH 7.0, 50 m*M* KCl, 1.0 m*M* DTT, and 1.0 m*M* MgCl_2_ and incubated at 25 °C for 30 min. The reaction was stopped by the addition of EDTA (2.0 m*M*). The reaction product ADP was quantified by analyzing the fluorescence signal of resorufin using a microplate spectrofluorometer Spectramax Gemini XPS (Molecular Devices) with λ ex = 540 nm and λ em = 590 nm. The kinetic constants were calculated using GraphPad Prism software (GraphPad Software).

### TNP-ATP kinase binding assay

Fluorescent measurements with an ATP analogue TNP-ATP (2’,3’-O-Trinitrophenyl-adenosine-5’-triphosphate, Jena Bioscience) were performed similar to previous study (36, 37). PKR^KD^ ^K296R/C551A^ (2.5 μ*M*) and PKR^KD^ ^K296R/C551A^-PK2 complex (2.5 μ*M*) were mixed with TNP-ATP (5 μ*M*) in the presence of various concentrations of ATP in a buffer consisting of 20 m*M* Tris pH 7.0, 50 m*M* KCl and 1.0 m*M* DTT, and were incubated at 25 °C for 30 min. The TNP-ATP fluorescence signals were measured using a microplate spectrofluorometer Spectramax Gemini XPS at λ ex = 408 nm and λ em = 538 nm. The kinetic constants were calculated using GraphPad Prism software (GraphPad Software).

## Data availability

The structural models and data in this study have been deposited in the RCSB Protein Data Bank under the following accession IDs: 9KV2 (PK2), 9KV3 (PK2ΔNTE), 9KV4 (PK2-BmHRI complex form I), and 9KV5 (PK2-BmHRI complex form II).

## Acknowledgements

We thank Dr. Penmetcha Kumar for discussions and critical reading of the manuscript. We thank Dr. Tomoyuki Numata for valuable discussions. We thank Dr. Takuya Miyakawa for information on the MBP-fusion protein expression system. We thank the beamline staff at Photon Factory (KEK, Tsukuba, Japan) in data collection. We also thank Natsuko Shirai for technical assistance. This work was performed with the approval of the Photon Factory Advisory Committee, Japan (Proposal No. 2016G091, 2018G110, 2023G156). This work was supported by JSPS KAKENHI (Grant-in-Aid for Young Scientists (B) 16K18512 and Grant-in-Aid for Scientific Research (C) 18K06103), Kato Memorial Bioscience Foundation, Agricultural Chemical Research Foundation, Koyanagi-foundation, and the natural science grant of the Mitsubishi Foundation.

## Author Contributions

D.T. designed the project. D.T. prepared samples for X-ray crystallographic studies and biochemical analysis. D.T. collected X-ray crystallographic data and analyzed X-ray crystallographic data. D.T. and Y.T. analyzed biochemical data. D.T. wrote the manuscript with assistance from Y.T.

## Extended Data Figure Legends

**Extended Data Figure 1.**
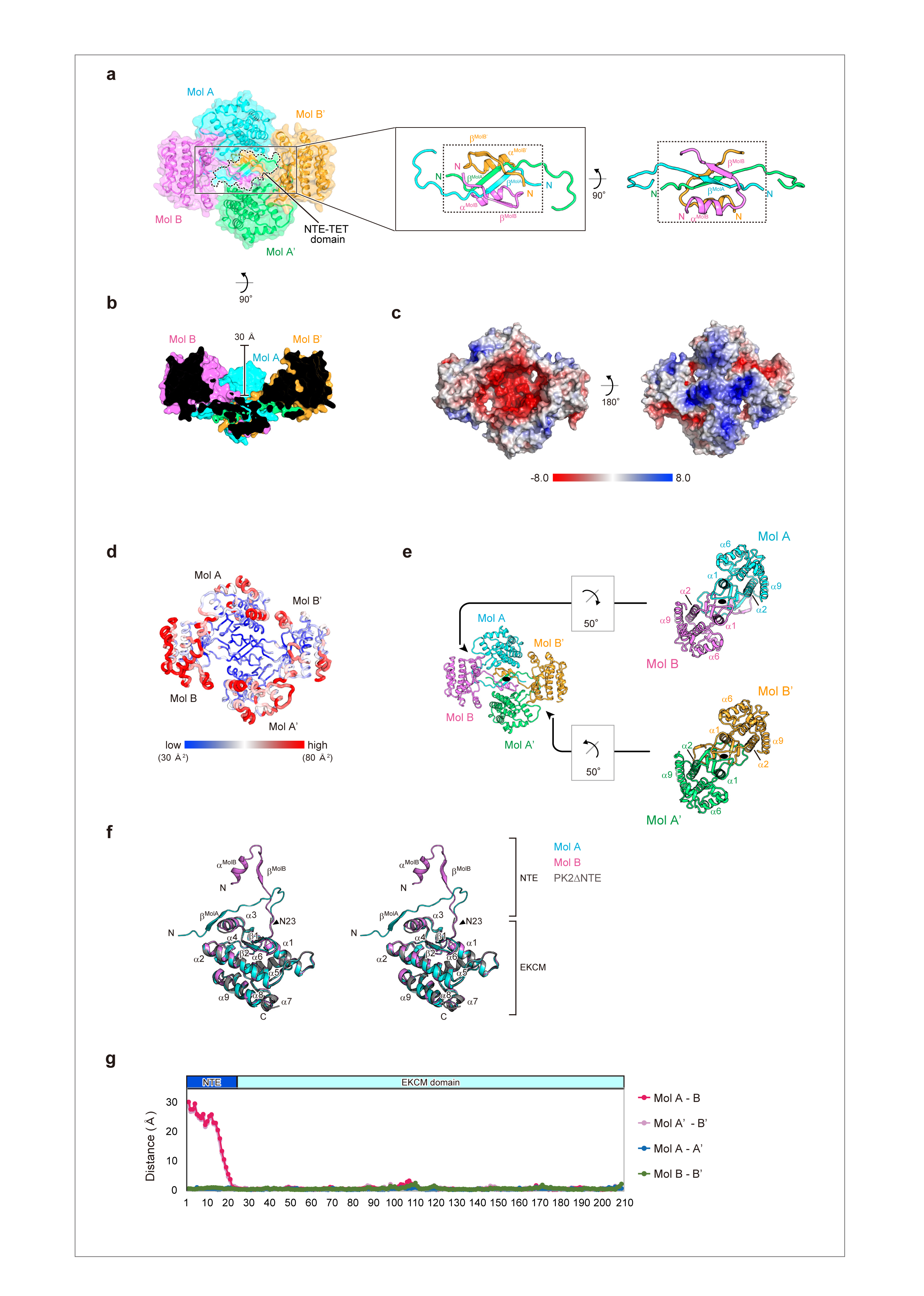
Structural analysis of PK2 homotetramer. **(a)** Transparent surface representation of PK2 homotetramer colored as in Fig. 1b (left panel). Close-up views of the NTE region at the center of PK2 homotetramer (middle and right panels). **(b)** A cross-section view of the structure of PK2 homotetramer, colored as in Fig. 1b but rotated 90° around the x-axis. **(c)** Electrostatic surface presentation of PK2 homotetramer colored red for negative, blue for positive, and white for apolar. **(d)** B-factor putty representation of PK2 homotetramer. Cα atoms with low temperature values are colored blue, while Cα atoms with high temperature values are colored red, as shown in the bar at the bottom. **(e)** Ribbon representation of PK2 homotetramer, and separated dimers are shown on the right. The two-fold symmetric axes are indicated by black ellipses. **(f)** Stereoview of PK2 protomers, colored as in Fig. 1d. **(g)** The distances between Cα atoms of PK2 promoters after superposition.

**Extended Data Figure 2.**
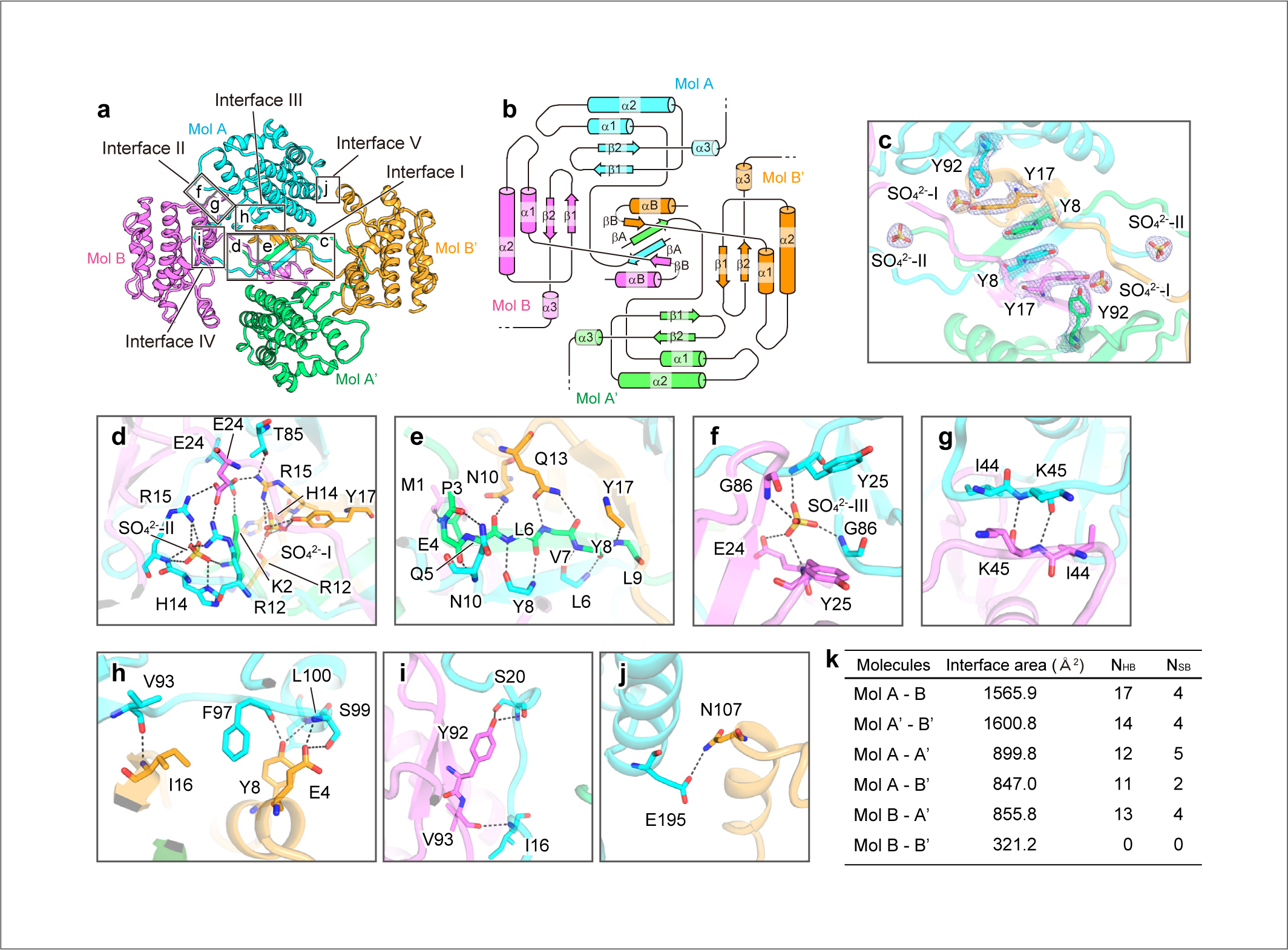
Interfaces of PK2 homotetramer. **(a)** The interfaces in the PK2 homotetramer are divided into five areas. Detailed interactions of each area are shown in c-j. **(b)** Topology representation of the PK2 homotetramer colored as same in Fig. 1b. For clarity, secondary structures from the N-terminus to the helix α3 of PK2 protomers are shown. **(c)** Six tyrosine residues form symmetric stacking interactions in the interface I. Four sulfate ions are symmetrically coordinated in the interface I. 2Fo-Fc map of the tyrosine residues and the sulfate ions are shown by blue mesh contoured at 2.0 σ level. **(d)** Two sulfate ions, SO_42-_-I and SO_42-_-II, are coordinated at the side of Mol B in the interface I. **(e)** Within the NTE-TET domain, β^MolA^, α^MolB^, and β^MolB^ of four protomers form hydrogen bonds. Side chains not involved in the interaction are omitted for clarity. **(f)** A sulfate ion SO_42-_-III is symmetrically coordinated in the interface II. **(g)** Main chains of Mol A and B form symmetric hydrogen-bonds in the interface II. **(h)** The EKCM domain of Mol A and NTE-TET domain form hydrogen bonds in the interface III. **(i)** The EKCM domain of Mol B and NTE-TET domain form hydrogen bonds in the interface IV. **(j)** A hydrogen bond is formed between Glu195 and Asn107 in the interface V. Note that the water-mediated hydrogen-bonding interactions are not shown, for clarity (d-j). **(k)** Summary of the total interface areas and numbers of hydrogen-bonds (N_HB_) and salt bridges (N_SB_) among the protomers of PK2 homotetramer.

**Extended Data Figure 3.**
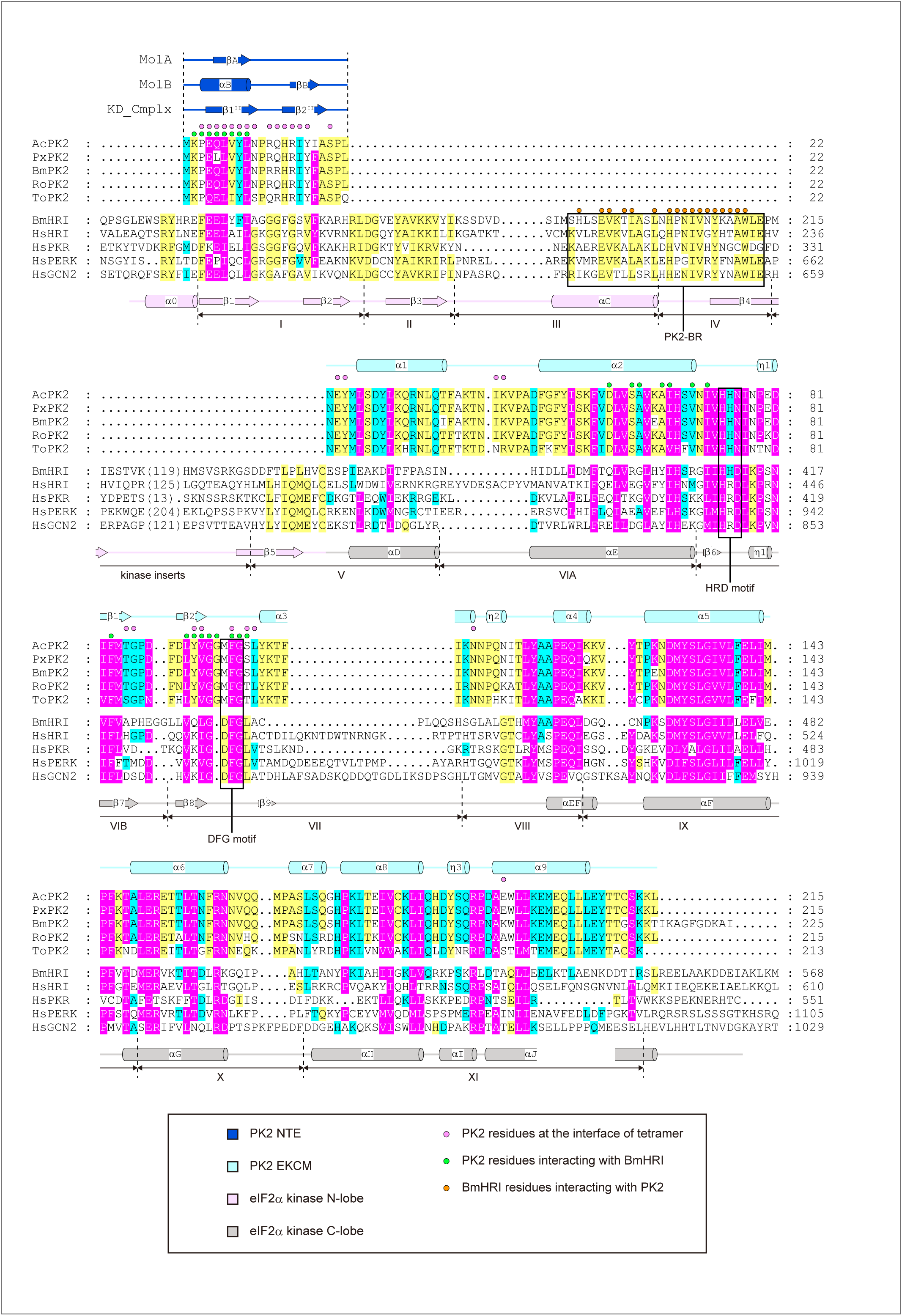
**Multiple sequence alignment of PK2 and eIF2**α **kinases.** Multiple sequence alignment of PK2 and eIF2α kinases, including PK2 from *Autographica californica* multiple nucleopolyhedrovirus (AcMNPV), *Plutella xylostella* multiple nucleopolyhedrovirus (PxMNPV), *Bombyx mori* nucleopolyhedrovirus (BmNPV), *Rachiplusia ou* MNPV (RoMNPV), and *Thysanoplusia orichalcea* NPV (ToNPV), and HRI kinases from *Bombyx mori* (Bm) and *Homo sapiens* (Hs), HsPKR, HsPERK, and HsGCN2. Conserved residues are shaded in magenta (80% conservation), cyan (60% conservation), and yellow (40% conservation). Pink circles indicate PK2 residues at the interface of homotetramer, and green circles indicate PK2 residues at the interface of the PK2-BmHRI complex. Orange circles indicate BmHRI residues at the interface of the PK2-BmHRI complex.

**Extended Data Figure 4.**
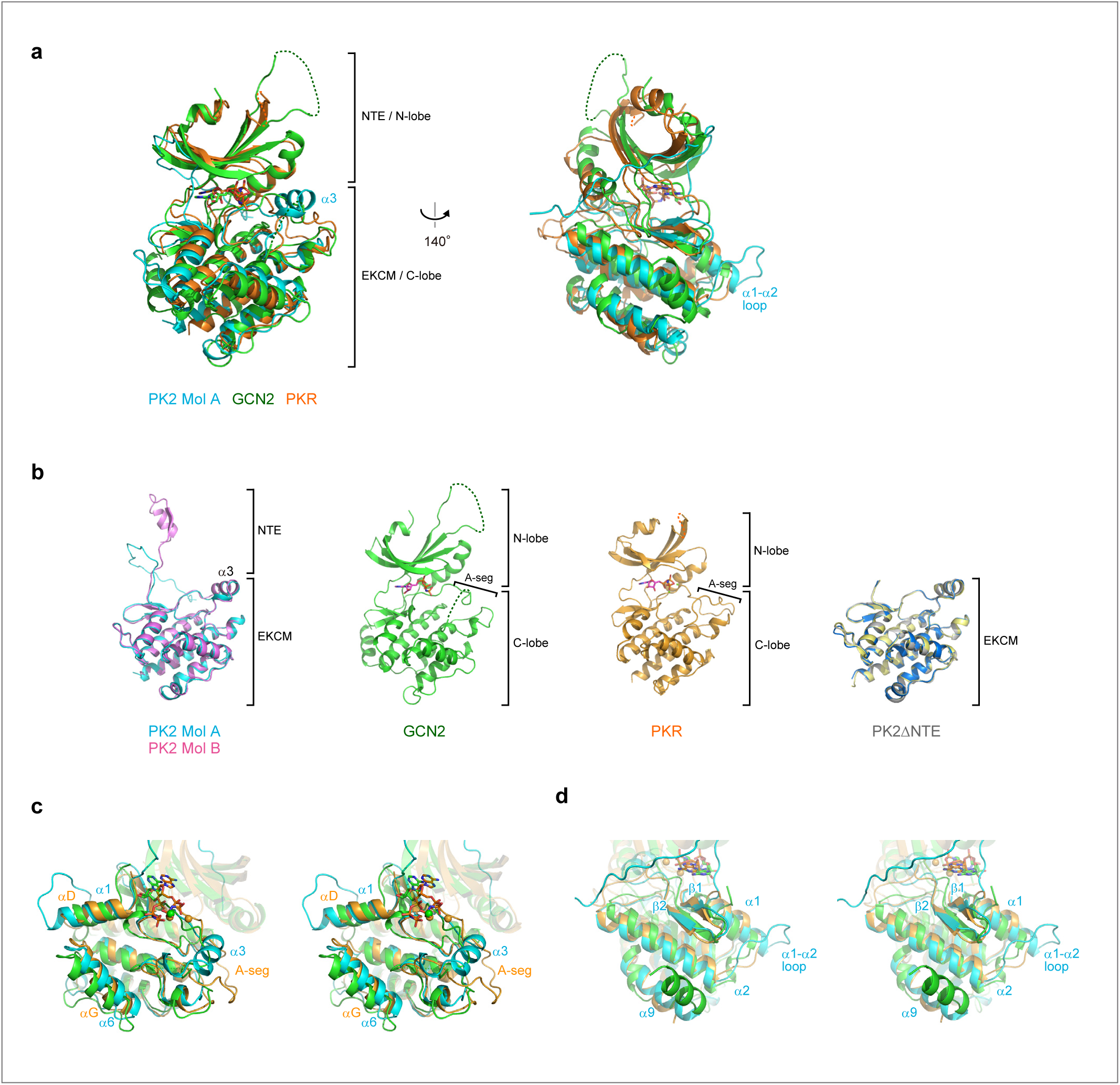
**Structural comparison of PK2 and eIF2**α **kinases. (a)** Superimposition of PK2 Mol A, and kinase domains of GCN2 (PDB ID: 1ZY5) and PKR (PDB ID: 2A19) colored cyan, green, and orange, respectively. **(b)** Superimposition of PK2 protomers, Mol A and Mol B, colored cyan and magenta, respectively. The kinase domains of GCN2 and PKR are also shown and colored green and orange, respectively. Superimposition of three molecules of PK2ΔNTE in the asymmetric unit colored gray, yellow, and blue. Views from the same direction as in the left panel. The activation segment (A-seg) and helix α3 are labeled. Dotted lines indicate flexible region in the crystal structures. **(c)** Close-up stereoview of superposition of PK2 protomer (Mol A), GCN2 and PKR colored cyan, green, orange, respectively. Compactly folded α3 of PK2 corresponds spatially to the activation segment (A-seg) of GCN2 and PKR. **(d)** Close-up stereoview of superposition of PK2 protomer (Mol A), GCN2 and PKR colored cyan, green, orange, respectively, as in Extended data Fig. 4c. The loop connecting α1 and α2 of PK2 forms an extended conformation compared to the loops of GCN2 and PKR.

**Extended Data Figure 5.**
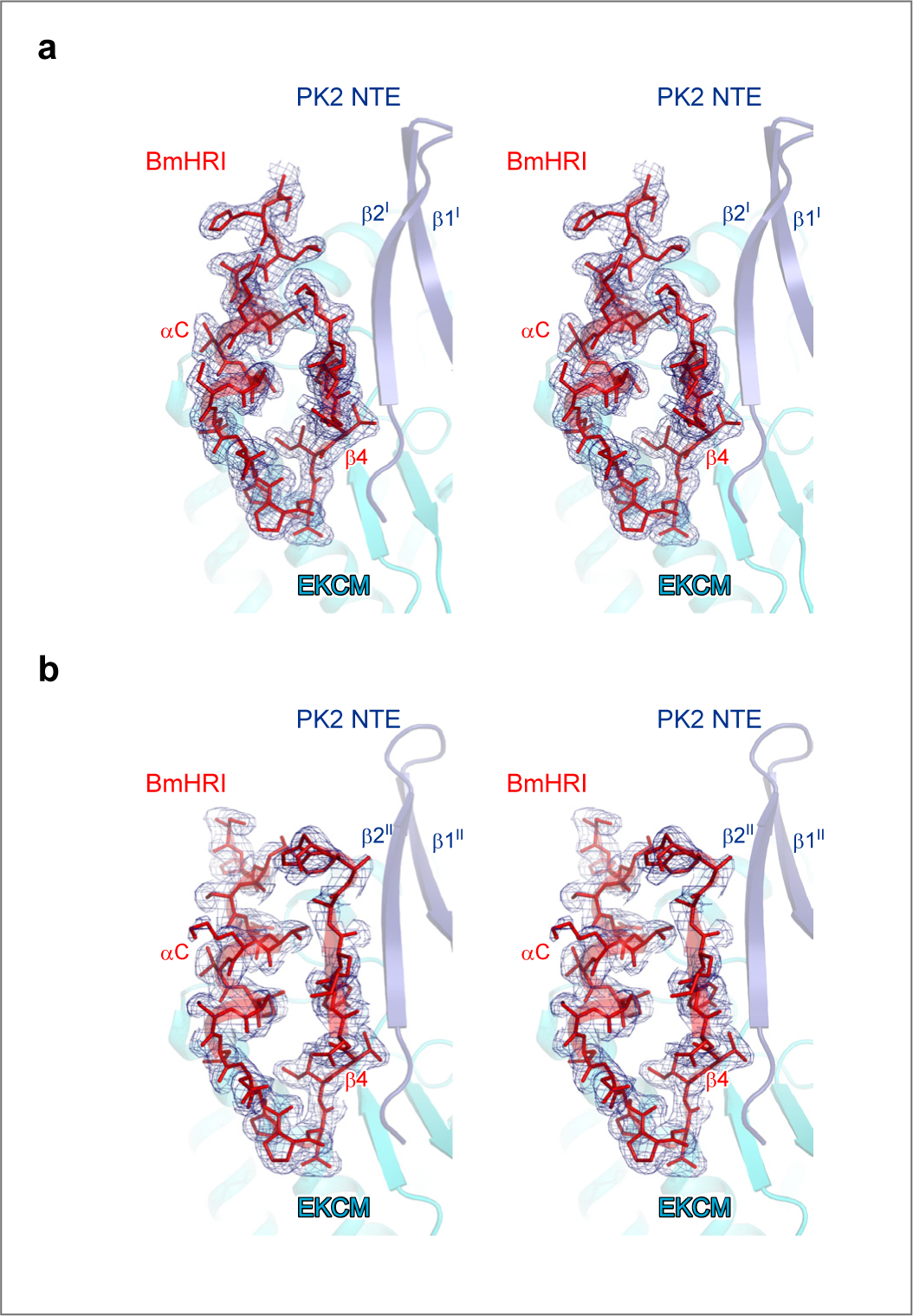
Stereoview of 2Fo-Fc electron density map for PK2-BR of BmHRI. **(a)** Stereoview of 2Fo-Fc map around BmHRI of form I complex contoured at 0.8σ. **(b)** Stereoview of 2Fo-Fc map around BmHRI of form II complex contoured at 0.8σ. Proteins are colored same as in Fig. 2c.

**Extended Data Figure 6.**
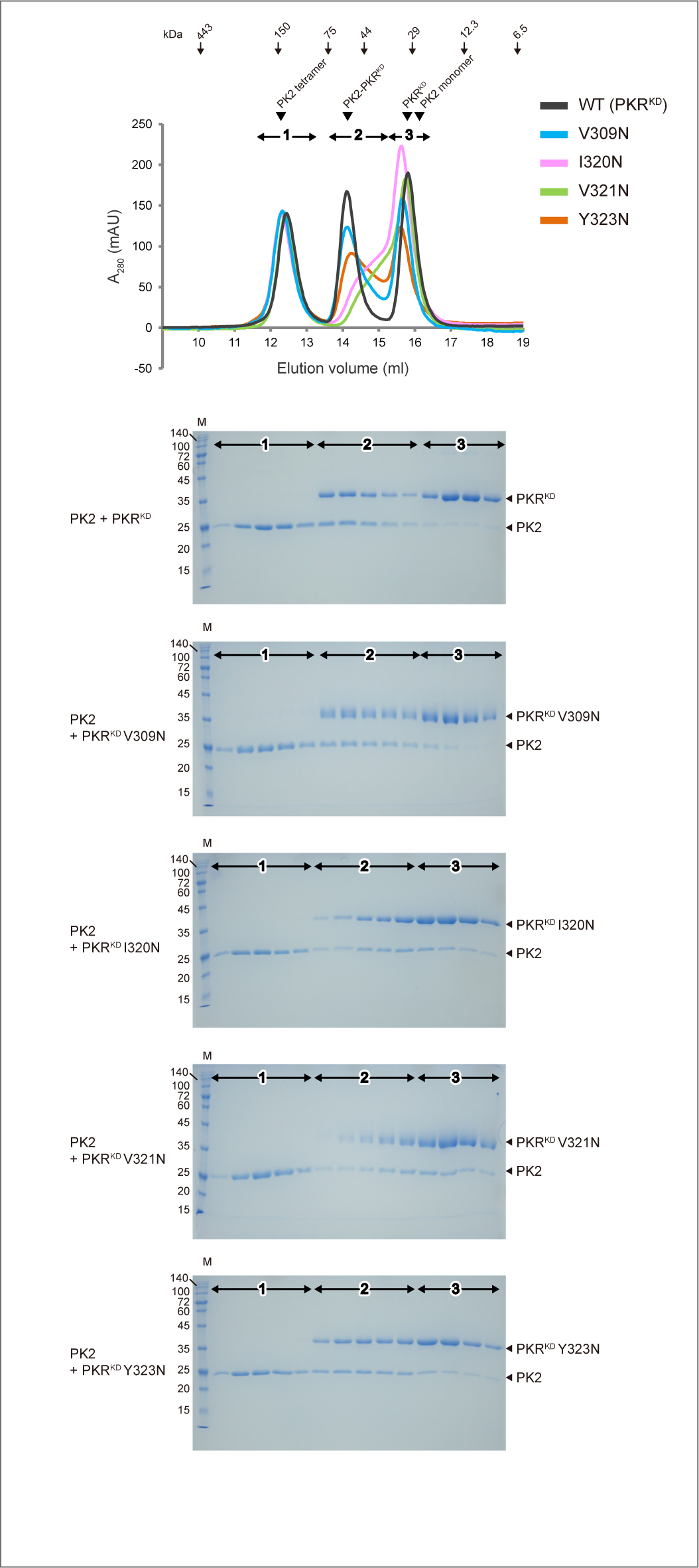
SDS-PAGE gels of fractions of the SEC analysis of the mixtures of PK2 and PKR_KD_ variants. Representative SDS-PAGE gels of the fractions of the SEC analysis of mixtures of PK2-PKR^KD^ complex, and PK2-PKR^KD^ mutant complexes, as presented in Fig. 4C. The mixtures are described on the left of the gels.

**Supplementary Table 1.**
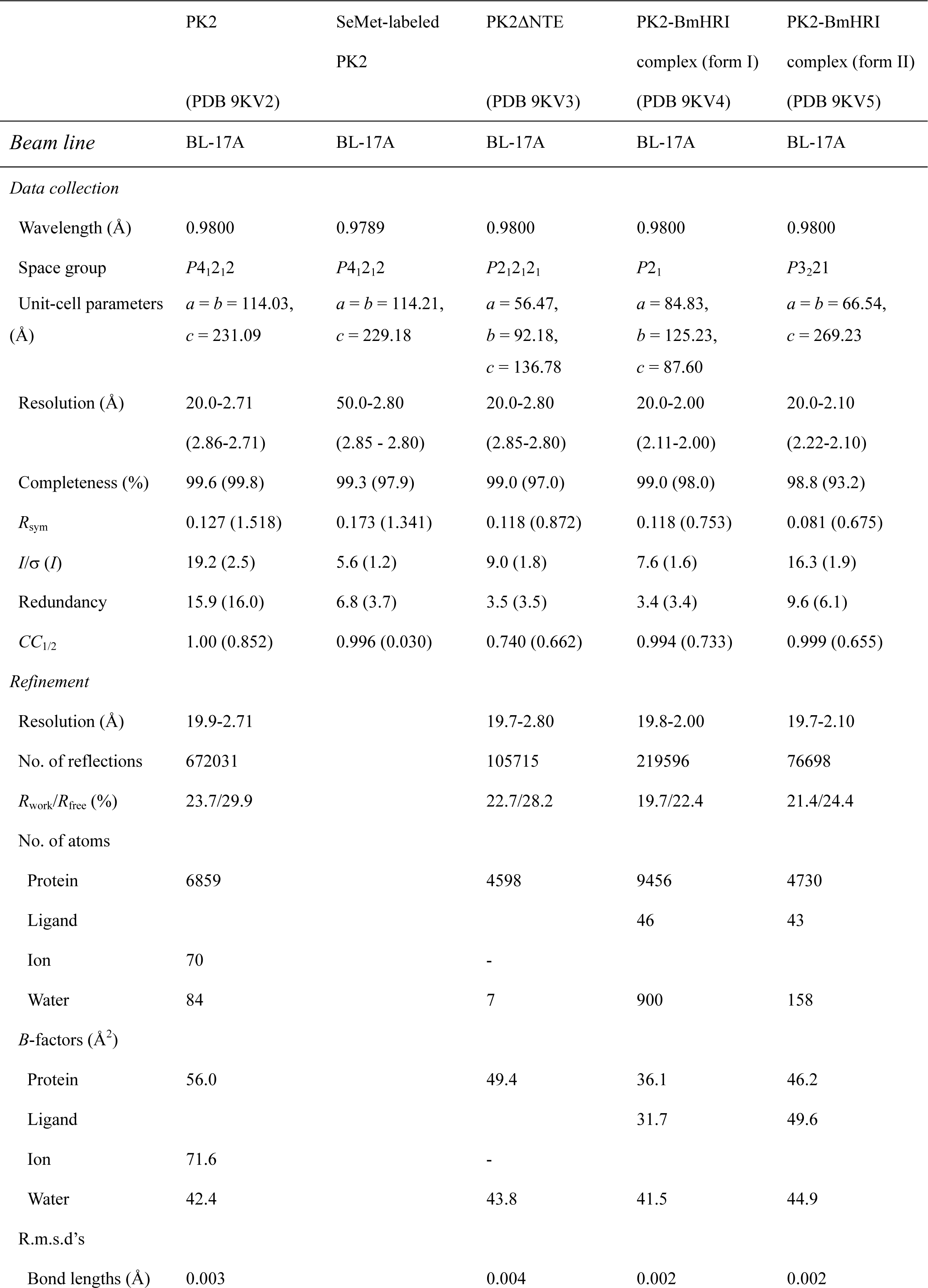
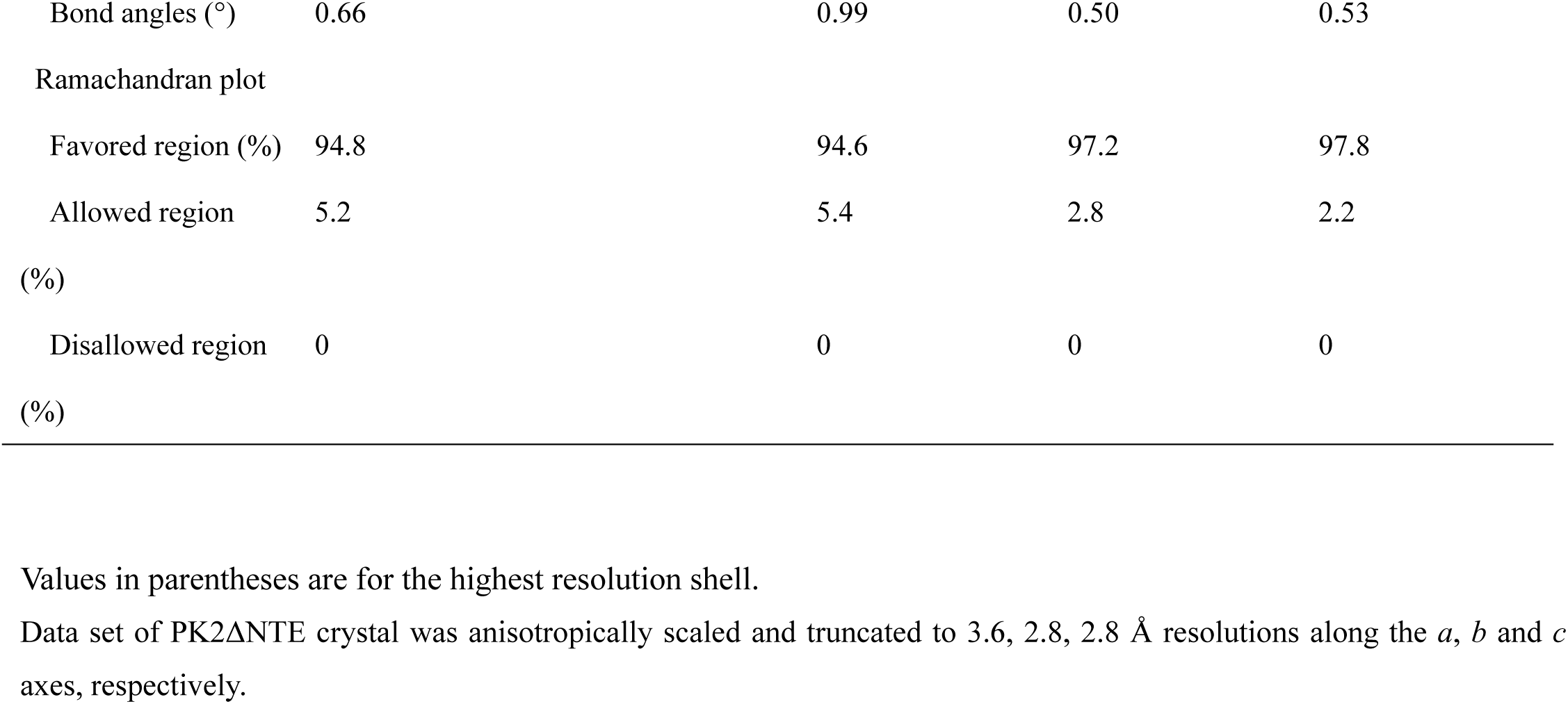
Data-collection statistics.

